# Cockayne syndrome patient iPSC-derived brain organoids and neurospheres show early transcriptional dysregulation of biological processes associated with brain development and metabolism

**DOI:** 10.1101/2023.10.17.562706

**Authors:** Leon-Phillip Szepanowski, Wasco Wruck, Julia Kapr, Andrea Rossi, Ellen Fritsche, Jean Krutmann, James Adjaye

## Abstract

Cockayne syndrome is a rare hereditary autosomal recessive disorder characterized by diverse neurological afflictions. However, little is known about the cerebral development in CS patients.

We generated neurospheres and cerebral organoids utilizing Cockayne Syndrome B Protein (CSB) deficient induced pluripotent stem cells derived from two patients with distinct severity levels of CS and healthy controls. The transcriptome of both developmental timepoints was explored using RNA-Seq and bioinformatic analysis to identify dysregulated biological processes common to both CS patients in comparison to control. CSB-deficient neurospheres displayed upregulation of VEGFA-VEGFR2 signaling pathway, Vesicle-Mediated transport and head development. CSB-deficient cerebral organoids exhibited downregulation of brain development, neuron projection development and synaptic signalling. We further identified upregulation of Steroid Biosynthesis as common to both timepoints, in particular upregulation of the Cholesterol Biosynthesis branch.

Our results provide insights into the neurodevelopmental dysregulation in CS patients and strengthen the theory, that CS is not only a neurodegenerative, but also a neurodevelopmental disorder.

## Introduction

Cockayne syndrome (CS), first described by Edward Alfred Cockayne in 1936^1^, is a rare hereditary autosomal recessive disorder characterized by severe photosensitivity, failure to thrive, cachectic dwarfism, segmental progeria, cataracts, and progressive multisystem degeneration. CS is also known for its various neurological afflictions such as mental retardation, sensorineural hearing loss, progressive microcephaly and hypomyelination.^2–4^

The two main mutated genes in CS patients are excision repair cross-complementing protein group 6 *ERCC6* (Cockayne Syndrome B Protein, CSB), which 2 of 3 patients exhibit a mutation in, and *ERCC8* (Cockayne Syndrome A Protein, CSA), which comprise the remaining 1/3.^2^

The CSB protein is a crucial part of the transcription-coupled nucleotide excision repair mechanism (TC-NER), where it detects the DNA lesion-stalled RNA Polymerase II (RNAPII) and initiates the repair process.^5^ However, while dysfunction of repair processes is associated with neurodegenerative diseases^6^, the loss of this repair mechanism cannot entirely explain the symptoms associated with CS patients.

Additionally to its function in the TC-NER, CSB has been implicated as enhancing the efficacy of the base excision repair and mitochondrial base excision repair mechanism^7–9^, is involved in the repair of double-strand breaks^10,11^ and has been implicated in enhancing inter-strand crosslinks repair.^12,13^ CSB has also been shown to be involved in transcription regulation. CSB deficient cells show up to 50% reduced transcription and dysregulation of thousands of genes.^14,15^

All these characteristics of CSB have been investigated employing mainly dermal fibroblasts of various CS patients, Chinese hamster ovary (CHO) and mouse model derived primary cells. Monotypic cell lines are typically well suited for initial gene function analyses and structure-function relationship studies but cannot fully illustrate the impact of a versatile protein such as CSB with multi-system manifestations. In 1997, the first mouse model of CS was described^16^, enabling investigation of primary cells of impacted tissues and whole-organisms studies. Since then, mouse models of all CS associated genes, *CSB, CSA, XPB, XPD* and *XPG* have been established.^17–21^ CS mouse models (except for *XPG*) generally exert a mild CS phenotype with reduced bodyweight, UV-sensitivity, mild neurodegenerative changes, and a surprising increase in skin tumorigenesis, a symptom not found in human CS patients. For modelling of severe CS, CSA- or CSB-deficient mice can be crossed with mice lacking in other genes of the TC-NER machinery, e.g., *XPA* or *XPC.*^18,22^

However, the discrepancy in the function of CS proteins and the need to combine two mutations to replicate specific characteristics emphasizes the need for humanized models. The advent of induced pluripotent stem cells (iPSCs) enabled investigators to develop reliable human disease models to study the effects of mutations on human tissues. Several neurodegenerative and mental disorders e.g., Alzheimer, Parkinsońs Disease, schizophrenia, and autism, have been the focal point of stem cell based neurological research. However, this rare disease has only been subject of few iPSC-based studies.^23–28^ Some of these studies concentrated on CS patient-derived dermal human fibroblasts or SH-SY5Y cells and only incorporated CSB-deficient iPSC lines for singular experiments. Still, these studies have provided great insight into the transcriptional dysregulation in CS patients. Several important pathways have been identified to be dysregulated in CS patient iPSC-derived neuronal networks e.g., axonogenesis, action potential of neurons, GH/IGF-1 signalling pathway, synaptic transmission, synaptogenesis and density of synapses.^26^ In accordance with these enrichment clusters, a reduced synapse density, a reduced spike number and an altered network synchrony has been identified in iPSC-derived neuronal networks.^26^ Two studies also identified dysregulation of miRNAs in CSB-deficient iPSCs and iPSC-derived neuronal networks.^23,26^ A successful rescue attempt via Crispr/Cas9 has demonstrated that restoration of one of two mutations in heterozygous mutants is sufficient to restore CSB function in CSB-deficient iPSCs, neural and mesenchymal stem cells.^27^ Also, pharmacological intervention in iPSC-derived neurons has shown that TRKB-agonists can increase arborization and neurite numbers in CSB-deficient iPSC-derived neurons.^25^

All these studies have used iPSCs or iPSC-derived 2D and 2.5D neuronal models. So far, no one has utilised the approach of self-organising organoids, which better recapitulate the cellular heterogeneity, structure, and functions of the primary tissues.

In this study, we sought to shed light on common transcriptional dysregulation associated with early cerebral development in CS patients. To this end, iPSC lines derived from (i) two individuals with CS and (ii) a healthy control were differentiated into expandable neural progenitor cells (NPCs) and cerebral organoids (COs). Next, we utilized Next-Generation Sequencing (NGS) to analyse RNA transcription at the NPC stage and after 60 days of CO differentiation. Here, we compared NPCs and COs generated from CS patient-derived iPSCs with each other and an unaffected control. Comparative gene expression analysis revealed dysregulation of several cellular pathways common between the two individuals. These included enrichment clusters which have been shown to be dysregulated in neurons, such as pathways associated with synaptogenesis, synaptic maintenance, axon guidance and p53 signalling pathway, but also pathways important for brain development and homeostasis, which have not been implicated in CS before. These included e.g., VEGFA-VEGFR2 signalling pathway, vesicle-mediated transport, and important metabolic pathways like steroid biosynthesis.

## Material and Methods

### iPSC Culture Methods

The iPSC lines derived from CS patients (CS789 and IUFi001) as well as the control line B4 used in this study have been previously described.^29–31^

One clone per iPSC line was used for further investigation. iPSC lines were cultured in mTeSR Plus medium (StemCell Technologies, Vancouver, Canada) supplemented with Penicillin/Streptomycin (P/S) (Gibco; Thermo Fisher Scientific, Inc., Waltham, MA, USA) on Matrigel-coated six-well plates (Corning, New York, NY, USA). The medium was changed every day and cells were passaged at 70%-80% confluency every 5-7 days using PBS^Ca-/Mg-^ (Life Technologies; Thermo Fisher Scientific, Inc., Waltham, MA, USA) supplemented with 0,5 mM UltraPure^TM^ EDTA (*Gibco*; Thermo Fisher Scientific, Inc., Waltham, MA, USA).

### Generation of Neurospheres

For the generation of neurospheres, iPSCs were differentiated using a modified version of a protocol previously described in our workgroup.^32–35^

Briefly, iPSCs were dissociated using TrypLE Express (*Gibco*; Thermo Fisher Scientific, Inc., Waltham, MA, USA) and seeded into ultra-low-attachment 96 well plates (Nunclon^TM^ Sphera^TM^, Thermo Fisher Scientific, Inc., Waltham, MA, USA) at 10000 cells/well with mTeSR medium supplemented with 10 µM Y-27632 (Tocris Bioscience; Biotechne, Minneapolis, USA). Neural induction was commenced at day 2 by replacing the medium with Neural Induction Medium (NIM; 47,5% Neurobasal A, 47,5% DMEM/F12, 2% B27 w/o Vitamin A, 1% N2 supplement, 1% GlutaMAX, 1% Penicillin/Streptomycin; all from Gibco) supplemented with 10 µM SB-431542 (Torics Bioscience), 5 µM Dorsomorphin (Tocris Bioscience) and 10 µM Y-27632. The medium was changed daily and at day 6, Y-27632 was omitted. At day 8, the spheroids were transferred to Poly-Ornithine and laminin-coated (Both Sigma-Aldrich; Merck KGaA, Darmstadt, Germany) six-well plates containing prewarmed neural differentiation medium (NDM; Neurobasal A, 2% B27, 1% N2, 1% Glut Amax, 1% P/S; all from Gibco) supplemented with 20 ng/mL of EGF and FGF2 (both PeproTech; Thermo Fisher Scientific, Inc., Waltham, MA, USA). The attached spheroids were further cultured in an incubator with 5% CO^2^ and 37°C with daily medium changes. At day 16 the neural rosettes were detached using STEMdiff^TM^ Neural Rosette Selection Reagent (StemCell Technologies, Vancouver, Canada) and transferred to non-adhesive 100 mm dishes containing NIM supplemented with 20 ng FGF2 and EGF. From day 16 onwards, neurospheres were cultured in a shaking incubator (New Brunswick S41i, Eppendorf) under continuous agitation (60 rpm) with 5% CO_2_ at 37°C with daily medium change.

### Generation of Organoids

For the generation of cerebral organoids, iPSCs were differentiated as described previously in our workgroup^32,34^, with minor modifications. In short, iPSCs were dissociated using TrypLE Express (Gibco; Thermo Fisher Scientific, Inc., Waltham, MA, USA) and seeded into ultra-low-attachment 96 well plates (Nunclon^TM^ Sphera^TM^, Thermo Fisher Scientific, Inc., Waltham, MA, USA) at 10000 cells/well with mTeSR medium supplemented with 10 µM Y-27632 (Tocris Bioscience; Biotechne, Minneapolis, USA). Neural induction was commenced at day 2 by replacing the medium with Neural Induction Medium (NIM; 47,5% Neurobasal A, 47,5% DMEM/F12, 2% B27 w/o Vitamin A, 1% N2 supplement, 1% GlutaMAX, 1% Penicillin/Streptomycin; all from Gibco) supplemented with 10 µM SB-431542 (Tocris Bioscience), 5 µM Dorsomorphin (Tocris Bioscience) and 10 µM Y-27632. The medium was changed daily and at day 6, Y-27632 was omitted. At day 8, the spheroids were transferred to non-adhesive 100 mm dishes containing Neural differentiation medium (NDM; Neurobasal A, 2% B27, 1% N2, 1% GlutaMAX, 1% P/S all from Gibco) supplemented with 20 ng/mL of EGF and FGF2 (both PeproTech; Thermo Fisher Scientific, Inc., Waltham, MA, USA). Organoids were further cultured in a shaking incubator (New Brunswick S41i, Eppendorf) under continuous agitation (60 rpm) with 5% CO_2_ at 37°C. The medium was changed daily until day 25. From day 25 to day 40, organoids were cultured in NDM supplemented with BDNF and NT-3 (both PeproTech; Thermo Fisher Scientific, Inc., Waltham, MA, USA). The medium was changed every 3 days. From day 40 onwards, the organoids were kept in NDM without further supplements with bi-weekly medium changes.

### Neurosphere/Organoid Sectioning and Immunocytochemistry

Cerebral organoids were fixed in ice-cold 4% paraformaldehyde (PFA) for 1 h at room temperature, washed three times with PBS and dehydrated for 24 h in 30% sucrose/PBS at 4°C.

Subsequently organoids were transferred to cryomolds containing Tissue-Tek OCT Compound 4583 embedding medium (Sakura Finetek, Umkirch, Germany), snap-frozen on liquid nitrogen and stored at −80°C. Organoids were cut into 10 µm sections using a Cryostat (CM1850, Leica, Nussloch, Germany) and captured on Superfrost Plus slides (Thermo Fisher Scientific, Inc., Waltham, MA, USA). For immunocytochemistry, cryosections were permeabilized with 0.5% Triton X-100 in PBS for 10 min and blocked with 3% BSA, 0,1% Triton X-100 in PBS for 1 h. Samples were then incubated overnight at 4 °C with the following primary antibodies: mouse anti-βIII-tubulin (1:400, Cell Signalling Technologies #4466S), rabbit anti-SOX2 (1:400, Cell Signalling Technologies #3579S), guinea pig anti-DCX (1:200, Merck Millipore #AB2253), mouse anti-KI67 (1:400, Cell Signalling Technologies #9449S), rabbit anti-cleaved CASP3 (1:400, Cell Signalling Technologies #9664S), rabbit Nestin (1:1000, Merck Sigma-Aldrich #N5413), mouse anti-MAP2 (Synaptic Systems, 1:500 #188011), guinea pig anti-NeuN (1:500, Synaptic Systems #266004), mouse anti-TAU (1:1000, Invitrogen, Thermo Fisher Scientific #MN1000), rabbit anti-S100B (1:100, Abcam #ab52642), goat anti SOX1 (1:200, R&D Systems, Biotechne #AF3369) and rabbit anti-gH2A.X (1:400, Cell Signalling Technologies #9718S). All antibodies are listed in Supplemental Table 1. Samples were washed three times with room-temperature (RT) PBS. Samples were further incubated with the appropriate secondary antibody conjugated with either Alexa-488, Alexa-555 or Alexa-647 (all 1:500, Invitrogen, Thermo Fisher Scientific) and the nuclear stain Hoechst 33258 (2 ug/mL, Sigma-Aldrich, Merck) for 2 h at RT. Slices were mounted with Fluoromount-G (Southern-Biotech, Birmingham, AL, USA) and dried overnight at room temperature. Fluorescent images were obtained using a LSM 700 microscope (Carl Zeiss, Jena, Germany), and processed using ZEN Blue software (Carl Zeiss) and ImageJ (National Institutes of Health, Bethesda, Maryland, USA).

### Image Analysis of Neurosphere and Organoid Histological Sections

Fluorescent images were obtained using a LSM 700 microscope (Carl Zeiss, Jena, Germany), and further analysed using ImageJ (National Institutes of Health, Bethesda, Maryland, USA).

To determine the number of Hoechst-, SOX2-, KI67- and gH2A.X-positive cells, 3-4 random sections were chosen per organoid/neurospheres, and the number of positive nuclei was determined by manual counting. For Cleaved-CASP3, ImageJ was used to determine the Integrated Density, area, and background fluorescence of 3-4 random sections per organoid/neurospheres. These values were then used to calculate the corrected total cell fluorescence (CTCF).

### Quantitative reverse transcription PCR (RT-qPCR)

Total RNA was extracted from iPSCs, 15-20 pooled day 30 neurospheres and 5-7 pooled day 60 organoids using TRIzol (Invitrogen; Thermo Fisher Scientific Inc., Waltham, MA, USA) and Direct-Zol RNA Mini Prep (Zymo Research, Freiburg, Germany). Extraction was performed according to the manufacturers protocol. 500 ng purified RNA was used for cDNA synthesis using a TaqMan reverse transcription reagent (Applied Biosystems; Thermo Fisher Scientific, Inc., Waltham, MA, USA).

Quantification of transcripts was performed by reverse transcription quantitative PCR (RT-qPCR) using the SYBR® Green RT-PCR assay (Applied Biosystems). Amplification, detection of specific gene products, and quantitative analysis were performed using the ViiA7 sequence detection system (Applied Biosystem). The expression levels were normalized relative to the expression of the housekeeping gene *RPLP0* and analysed via the 2^−ΔΔ-CT^ method. RT-qPCR data are depicted as mean values. Primers are listed in Supplementary Table S1.

### Next-Generation Sequencing

1µg RNA of each sample was subjected to bulk Next-Generation Sequencing (NGS) at BGI Genomics (BGI Group, Shenzhen, Guangdong, China). BGI performed the RNAseq (Transcriptome) library preparation via stranded mRNA enrichment method. Sequencing was performed on a DNBSeq PE100 platform with 20M clean reads per sample.

FPKM (Fragments Per Kilobase per Million mapped fragments) and count values aligned by the BGI via the HISAT software^36^ to the reference genome GCF_000001405.38_GRCh38.p12 were imported into the R/Bioconductor environment.^37^ Genes were considered expressed when the counts value was greater than the threshold of t=5. Venn diagrams were generated based on expressed genes using the R package VennDiagram.^38^ Dendrograms and heatmap were generated via the R methods “heatmap.2” and “hclust” from the R package gplots^39^ or the R core functionality using Pearson correlation as similarity measure. Gene ontologies (GOs) were analyzed via the GOstats R package.^40^ KEGG pathways^41–44^ were downloaded from the KEGG database in July 2020 and tested for over-representation of genes via the R-built-in hypergeometric test. Dot plots of the most significantly over-represented pathways or GOs were generated via the R package ggplot2.^45^ Differentially expressed genes between two conditions were determined via the R-built-in Fisher’s exact test using the criteria p < 0.05 and the ratio > 1.5 for up- or ratio < 2/3 for down-regulation.

Comprehensive functional analysis of the clustered GO biological processes and pathways (KEGG pathways, Reactome Gene Sets, Canonical pathways, and CORUM) of gene-sets based on previous Venn analysis was performed using Metascape (http://metascape.org, accessed on 27 July 2022 and 03 March 2023).^46^

To identify developmental age and enriched brain regions in the neurospheres and organoids, NGS datasets associated with fetal brain regions were downloaded from the Allen Brain Atlas (ABA) (https://www.brainspan.org/, accessed on 19.01.2022).^47^ Genes specific for brain regions were extracted from the ABA expression RPKM (reads Per Kilobase per Million mapped reads) data by first filtering only genes with a coefficient of variation (cv) greater than the median cv. Thereafter for each brain region the genes greater than the threshold t95 at the mean of the 95-quantiles of all samples were extracted. Heatmaps of enrichment in the ABA were made via the R package GSVA^48^ (gene set variation analysis) and represent the GSVA enrichment scores of the dedicated brain regions. Brain regions are abbreviated by the acronyms: occipital neocortex—Ocx, primary motor-sensory cortex (samples)—M1C-S1C, amygdaloid complex—AMY, medial ganglionic eminence—MGE, posterior (caudal) superior temporal cortex (area 22c)—STC, upper (rostral) rhombic lip—URL, caudal ganglionic eminence—CGE, dorsal thalamus— DTH, anterior (rostral) cingulate (medial prefrontal) cortex—MFC, dorsolateral prefrontal cortex—DFC, orbital frontal cortex—OFC, lateral ganglionic eminence—LGE, inferolateral temporal cortex (area TEv, area 20)—ITC, hippocampus (hippocampal formation)—HIP, ventrolateral prefrontal cortex—VFC, parietal neocortex—PCx.

### Western Blotting

To harvest total protein, 10-15 neurospheres and 4-6 organoids per cell line were rinsed twice with PBS and then lysed in RIPA buffer (Sigma-Aldrich; Merck KGaA, Darmstadt, Germany) supplemented with Protease Inhibitor Cocktail and Phosphatase Inhibitor Cocktail (Roche, Mannheim, Germany). Protein content of lysates was quantified using the Pierce^TM^ Protein Assay Kit (Thermo Fisher Scientific Inc., Waltham, MA, USA). 20 µg of the protein samples were then heat-denatured and subsequently separated on a 4-12% Bis-Tris gel (Invitrogen; Thermo Fisher Scientific). Separated proteins were then transferred to a 0.45 µm nitrocellulose membrane via wet blotting for 3h at 300 mA. Membranes were blocked in Tris-buffered saline/0.1% Tween-20 containing 5% skim milk and afterwards stained with the following antibodies: rabbit anti-CSB (1:1000, GeneTex Inc. #GTX104589), rabbit anti-SQLE (1:1000, Proteintech #12544-1-AP), mouse anti-p53 (1:1000, Merck OP43) and mouse anti β-actin (1:4000, Cell Signalling Technologies #4967S). All antibodies are listed in Supplemental Table 1. After incubation, blots were washed three times with Tris-buffered saline/0.1% Tween-20 and subsequently stained with appropriate secondary antibodies. Following three washing steps with Tris-buffered saline/0.1% Tween-20, the protein bands were visualized with ECL Western Blotting Detection Reagents (Cytiva, Freiburg, Germany) in a UV chamber. The resulting bands were quantified using grey scale analysis with ImageJ (National Institutes of Health, Bethesda, Maryland, USA).

## Results

### Both control and CSB-iPSCs efficiently differentiate into neural progenitors

For this project, we utilized two previously established iPSC lines generated from a patient with cerebro-oculo-fascio-skeletal syndrome (CS789), a patient with classical CS (IUFi001), as well as the established control line B4. The CS789 line carries a homozygous p.R683X mutation in exon 10 of the *ERCC6* gene, leading to a stop codon and a predicted truncated protein. The IUFi001 line carries a compound mutation, p.K377X in exon 5 and p.R857X in exon 15 of the *ERCC6* gene (Supplementary Figure S1).^29–31^

To investigate how CS affects early human neurodevelopment, we implemented our published suspension protocols for neurospheres (NS) (Fig. 1A) and cerebral organoids (COs) (Fig. 2A).^32,34^ The neurospheres were cultured for 30 days.

**Fig. 1:**
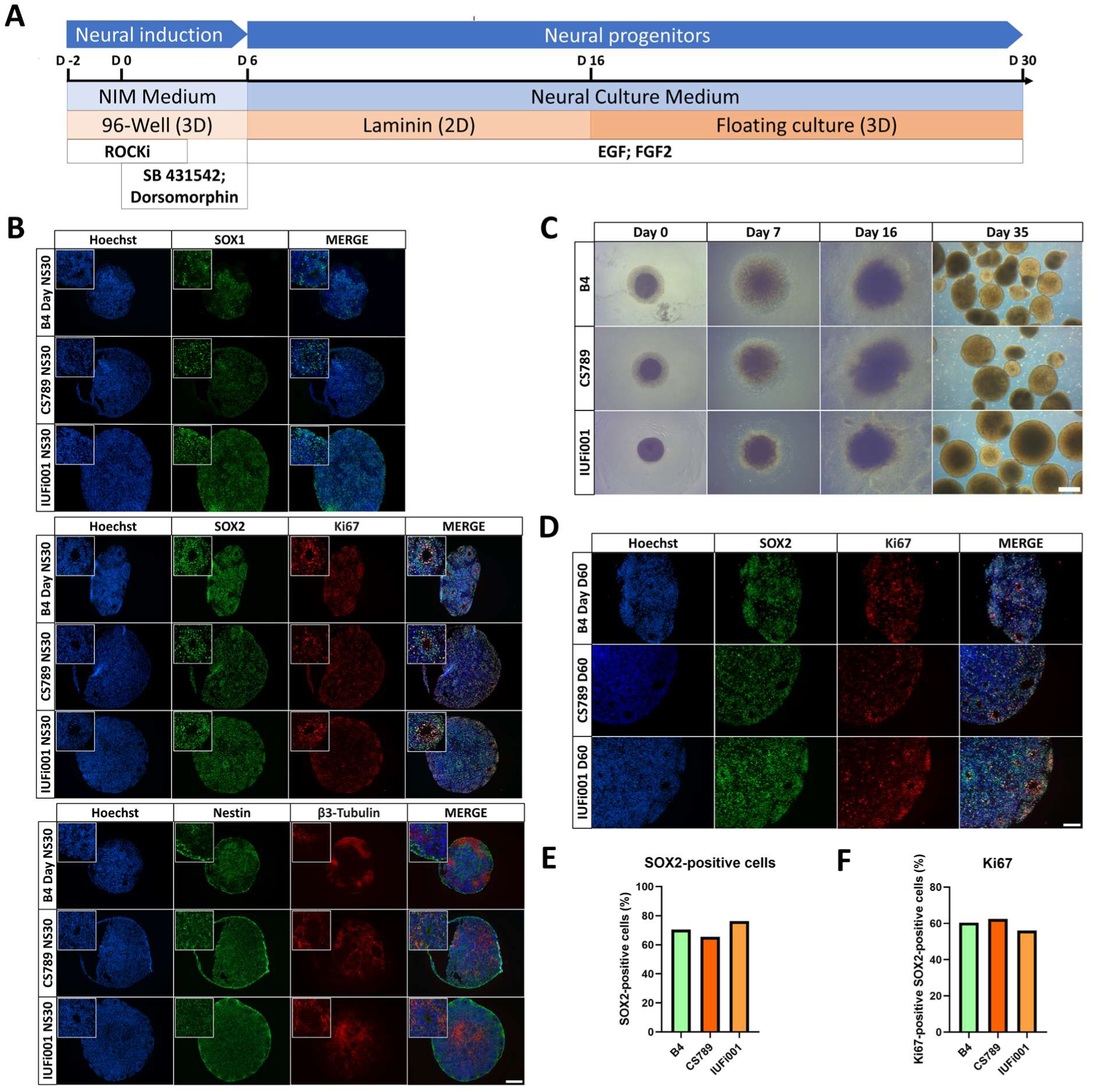
Generation and characterization of CS neurospheres. (**A**) Schematic outline of the protocol to generate iPSC-derived neurospheres. (**B**) Representative immunocytochemistry images of the distribution of cells expressing SOX1, SOX2, Ki67, Nestin and β3-Tubulin. 100x magnification, scale bar 200 µm. (**C**) Representative brightfield images of control and CS neurospheres at day 0, 7, 16 and 35 of differentiation. Scale bar, 200 µm. (**D**) Representative immunocytochemistry images of the distribution of cells expressing SOX2 and Ki67. 200x magnification, scale bar 100 µm. (**E**) Quantification of the SOX2-positive, Hoechst-positive cells in CTRL (B4), CS789 and IUFi001 neurospheres. (**F**) Quantification of the Ki67-positive cells within the SOX2-positive cells in CTRL (B4), CS789 and IUFi001 neurospheres. (**E,F**) Approximately ten random fields from three distinct neurospheres were analysed.

**Fig. 2:**
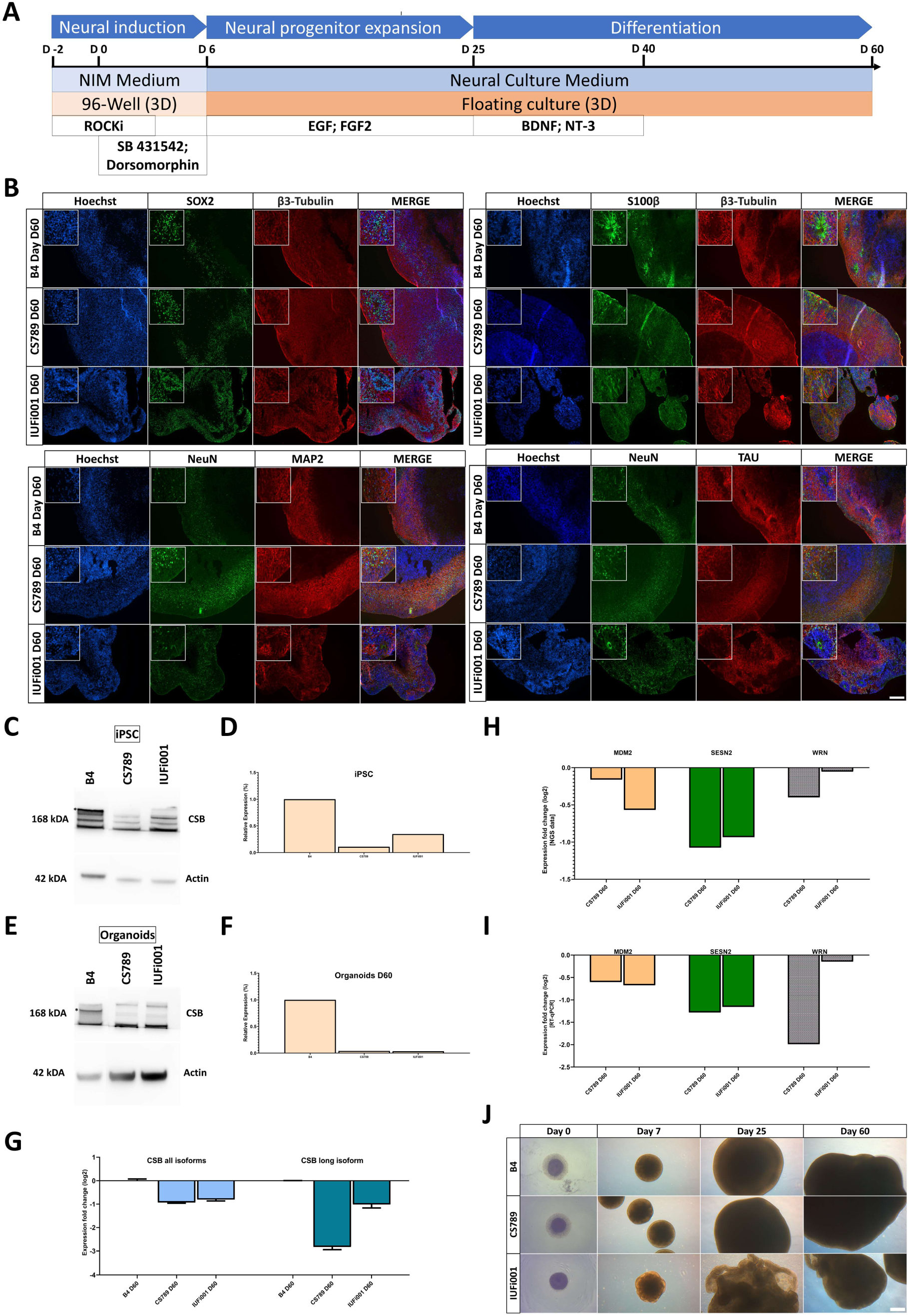
Generation and characterization of CS organoids. (**A**) Schematic outline of the protocol to generate iPSC-derived cerebral organoids. (**B**) Representative immunocytochemistry images of the distribution of cells expressing SOX2, β3-Tubulin, S100, NeuN, MAP2 and TAU. 100x magnification, scale bar 200 µm. (**C**) Western blot analysis for full length CSB at the iPSC stage. (**D**) Quantification of CSB Western blot analysis at the iPSC stage in CTRL (B4), CS789, and IUFi001. CSB expression of CS789 and IUFi001 is compared to CSB expression in CTRL (B4) (**E**). Western blot analysis for full length CSB at day 60 of organoid differentiation. (**F**) Quantification of CSB western blot analysis at iPSC stage in CTRL (B), CS789, and IUFi001. CSB expression of CS789 and IUFi001 is compared to CSB expression in CTRL (B4). (**G**) qRT-PCR analysis of full length CSB mRNA expression and all isoform CSB mRNA expression in CS789 and IUFi001 organoids relative to control organoids. (**H**) Relative mRNA expression analysis of DNA damage-related genes *MDM2*, *SESN2* and *WRN* in CS789 and IUFi001 organoids compared to CTRL (B4). (**I**) qRT-PCR analysis of *MDM2*, *SESN2* and *WRN* mRNA expression in CS789 and IUFi001 organoids relative to CTRL (B4). (**J**) Representative brightfield images of control and CS organoids at day 0, 7, 25 and 60 of differentiation. Scale bar 200 µm.

By day 30 of differentiation, immunocytochemical analysis revealed that control and CS NS were composed mostly of SRY-Box Transcription Factor 2 (SOX2)^+^, SRY-Box Transcription Factor 1 (SOX1)^+^ and Nestin^+^ neural progenitor cells (NPCs) organized in ventricular zone (VZ)-like structures surrounded by pan-neuronal marker Tubulin Beta 3 Class III (TUBB3)^+^ early-borne neurons (Fig. 1B).

During differentiation, all cell lines behaved similarly with regards to attachment, formation of rosettes and generation of NS (Fig. 1C). Manual assessment of SOX2^+^ nuclei (Fig. 1E) and SOX2 and Marker of Proliferation Ki-67 (Ki-67) double-positive nuclei (Fig. 1F) did not reveal differences in the number of SOX2^+^ NPCs and proliferating SOX2^+^ NPCs between all cell lines. Further ICC analysis can be found in Supplementary Figure S2.

By comparing the B4, CS789 and IUFi001 NS RNA-Seq datasets with transcriptional datasets from the Allen Brain Atlas we attempted to determine the developmental age of the NS (Supplementary Figure S3) as described previously.^32,49^ We observed that the transcriptomes of our day 30 NS are equivalent to 8-9 weeks post-conception fetal brains (Supplementary Figure S3).

These results imply, that control and CS patient-derived iPSCs can be efficiently differentiated into NS following our protocol. Patient-derived NS do not seem to exhibit changes in cyto-architecture, cellular makeup or proliferation in comparison to control.

### Both control and CSB-iPSCs efficiently differentiate into cerebral organoids

To elucidate transcriptional regulation at advanced stages of early human neurodevelopment in CS patients, we also cultured cerebral organoids for 60 days.

By day 60 of differentiation, immunocytochemical staining revealed that both CS and control lines share a common cyto-architecture. The outer layer (∼300-500 nm) of these COs is composed of SOX2^+^ NPCs self-organized into VZ-like structures surrounded by early-borne and mature neurons.

The mature neurons are Microtubule Associated Protein 2 (MAP2)^+^, Microtubule Associated Protein Tau (TAU)^+^ and RNA Binding Fox-1 Homolog 3 (NeuN)^+^ in COs of all cell lines (Fig. 2B). All neurons including the early-borne neurons are stained with TUBB3. Staining of S100 Calcium Binding Protein B (S100B) reveals the presence of radial glial cells and/or early astrocyte progenitors in COs of all three cell lines. Further ICC analysis can be found in Supplementary Figure S4.

We performed a Western blot to detect CSB at the iPSC and day 60 CO stage (Fig. 2C, E). This revealed markedly reduced levels of CSB in both patient-derived iPSC lines and COs in comparison to control (Fig. 2D, F). The full Western Blot can be found in Supplementary Figure S5. RT-qPCR revealed a ∼50% translation reduction of all CSB isoforms in both patient-derived cell lines, a ∼90% translation reduction of the full-length isoform in the CS789 line and a ∼50% reduction in the IUFi001 line (Fig. 2G).

Prior to NGS analysis, we performed RT-qPCR for the genes MDM2 Proto-Oncogene, E3 Ubiquitin Protein Ligase (MDM2), Sestrin-2 (SESN2) and Werner Syndrome RecQ Like Helicase (WRN), which were expected to be regulated in our patient-derived COs (Fig. 2I). The dysregulation was later confirmed by the NGS data (Fig. 2H).

Over the course of the CO differentiation, control and CS789 cell lines behaved similarly with respect to growth. The IUFi001 COs flattened and folded early in the differentiation and were broken up into smaller organoids by the shear forces of the spinning incubator (Fig. 2J). Still, except for the size, the cyto-architecture of the IUFi001 COs is comparable to the control and CS789 cell line.

By comparing the CO RNA-Seq datasets with datasets from the Allen Brain Atlas we attempted to determine the developmental age of the COs (Supplementary Figure S3). We observed that the transcriptomes of the day 60 COs are approximate to 13-21 weeks post-conception fetal brains (Supplementary Figure S3).

Collectively, these results suggest that control and CS patient-derived iPSCs can efficiently differentiate into COs following our protocol. Patient-derived CS COs do not seem to exhibit major changes in cyto-architecture in comparison to control and recapitulate aspects of early human neurodevelopment, but present expected dysregulation of damage-related gene expression.

### CSB-neurospheres of patients with different severity show distinct but partially overlapping transcriptome dysregulation

To gain insight into the early neurodevelopmental transcriptional differences between CS patients and healthy individuals, we performed transcriptome analysis of the day 30 NS via RNA-Seq. Hierarchical clustering (Supplementary Figure S6) and principal component analysis (Supplementary Figure S6) of all datasets (iPSCs, neurospheres and organoids of B4, CS789 and IUFi001) revealed correct clustering of the respective timepoints to each other with no outliers.

Next, we compared the transcriptome of CS789 NS to control NS. We found that 641 genes are exclusively expressed in CS789 NS, 635 genes exclusively expressed in control NS, and 14001 genes expressed in both. Of these commonly expressed 14,001 genes, 2302 genes are upregulated, and 430 genes downregulated in CS789 NS in comparison to control (Fig. 3A). With these identified differentially expressed genes (DEGs) we performed enrichment analysis to identify the most dysregulated Kyoto Encyclopaedia of Genes and Genomes (KEGG) pathways in CS789 NS. The three most upregulated KEGG pathways are - *Protein processing in endoplasmatic reticulum, Adherens junction* and *Hippo signalling pathway* (Fig. 3C). The five most downregulated KEGG pathways are *Ribosome, Oxidative Phosphorylation* and *Thermogenesis* (Fig. 3D).

**Fig. 3:**
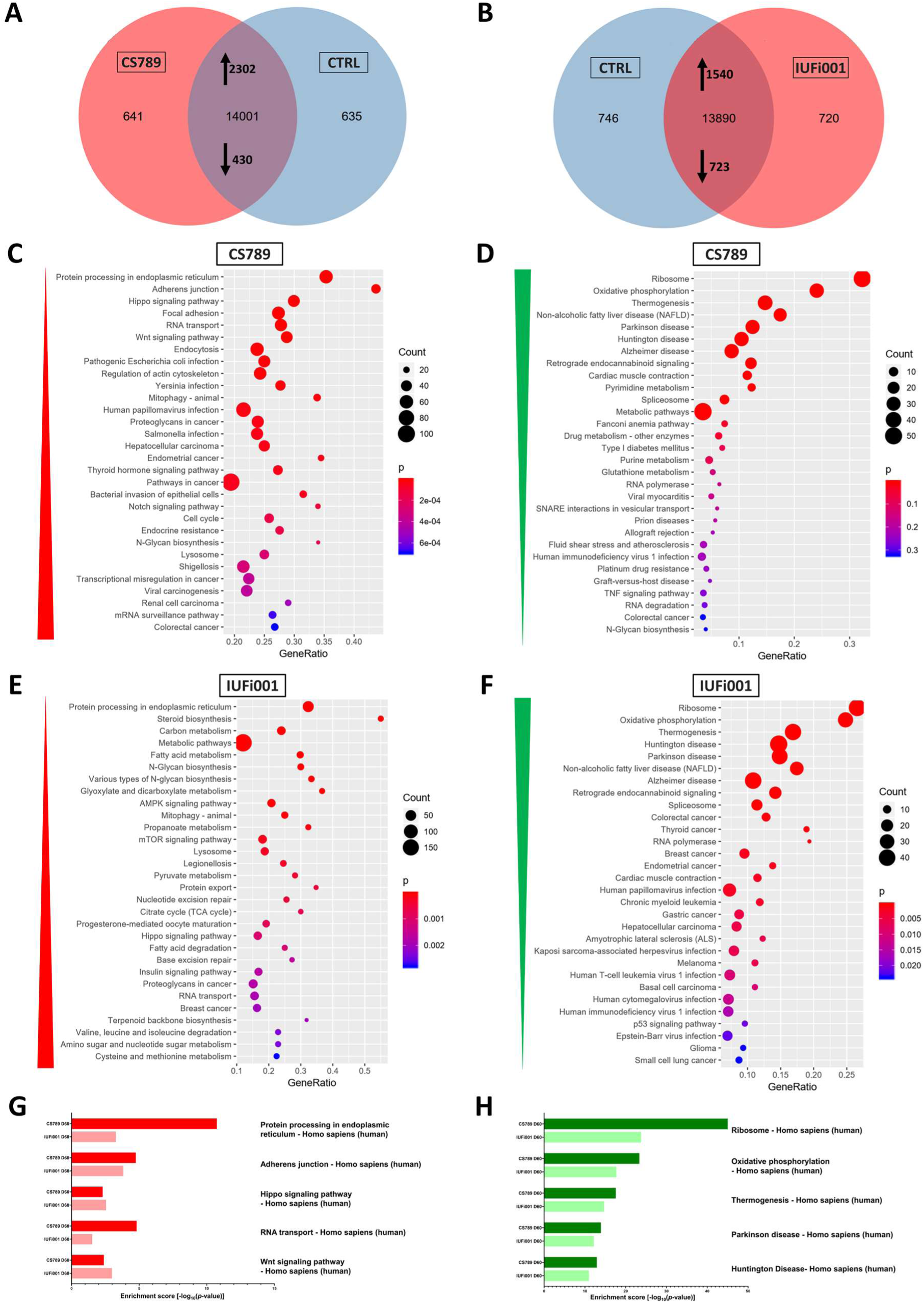
Global transcriptome and associated pathway analysis of control and CS neurospheres at day 30. (**A**) Venn diagram showing genes expressed only in CS789 neurospheres (641), in CTRL (B4) neurospheres (635) and common to both (14001) (detection p value < 0.05). (**B**) Venn diagram showing genes expressed only in IUFi001 neurospheres (720), in CTRL (B4) neurospheres (746) and common to both (13890) (detection p value < 0.05). (**C,D**) Dot plots showing the Top 30 differentially regulated KEGG pathways (C) in the 2302 significantly upregulated DEGs in day 30 CS789 neurospheres in comparison to CTRL (B4) (D) and in the 430 significantly downregulated DEGs in day 30 CS789 neurospheres in comparison to CTRL (B4). (**E,F**) Dot plots showing the Top 30 differentially regulated KEGG pathways (E) in the 1540 significantly upregulated DEGs in day 30 IUFi001 neurospheres in comparison to CTRL (B4) (F) and in the 723 significantly downregulated DEGs in day 30 IUFi001 neurospheres in comparison to CTRL (B4). (**G**) Bar chart of the differentially upregulated KEGG pathways (Top 5 ranked) common between day 30 CS789 and IUFi001 neurospheres in comparison to CTRL (B4) neurospheres. (**H**) Bar chart of the differentially downregulated KEGG pathways (Top 5 ranked) common between day 30 CS789 and IUFi001 neurospheres in comparison to CTRL (B4) neurospheres.

Subsequently, the transcriptome of the IUFi001 NS was compared to control NS. We identified 720 genes to be exclusively expressed in IUFi001 NS, 746 genes exclusively expressed in control NS, and 13,890 genes expressed in both. Of the commonly expressed 13,890 genes, 1540 genes are upregulated, and 723 genes downregulated in IUFi001 NS in comparison to control (Fig. 3B). The three most upregulated KEGG pathways in IUFi001 NS are - *Protein processing in endoplasmatic reticulum, Steroid biosynthesis* and *carbon metabolism* (Fig. 3E). The three most downregulated KEGG pathways are *Ribosome, Oxidative Phosphorylation* and *Thermogenesis* (Fig. 3F). Since we want to identify dysregulation common to all types of CS, we compared the 50 highest up- and downregulated KEGG pathways in both patient-derived NS in comparison to control. The five most upregulated KEGG pathways common in CS789 and IUFi001 neurospheres are *Protein processing in endoplasmatic reticulum, Adherens junction, Hippo signalling pathway, RNA transport and Wnt signalling pathway* (Fig. 3G). The five most downregulated KEGG pathways common in CS789 and IUFi001 neurospheres are *Ribosome, Oxidative Phosphorylation, Thermogenesis, Parkinson disease* and *Huntington Disease* (Fig. 3H). The complete list of pathways common in the 50 most severely dysregulated KEGG pathways can be found in Supplemental Table 2.

These results indicate that while there are ample differences in gene expression between distinct individuals and severity grades of CS, there is also common dysregulation. And while most symptoms manifest only postnatally in all but the most severe form of CS, this common dysregulation can already be detected in carriers of less severe forms at the NPC stage.

### CSB-organoids of patients with different severity show distinct but partially overlapping transcriptome dysregulation

To investigate the molecular differences between CS patients and healthy individuals at a more advanced stage of brain development (BD), we performed transcriptome analysis of the day 60 cerebral organoids.

We first compared the transcriptome of the CS789 COs to control COs. We identified 443 genes as exclusively expressed in CS789 COs, 1006 genes exclusively expressed in control COs, and 14030 genes expressed in both. Of these commonly expressed 14030 genes, 373 genes are upregulated, and 2163 downregulated in CS789 COs in comparison to control (Fig. 4A). We utilized the DEGs to perform enrichment analysis to identify the most dysregulated KEGG pathways in CS789 COs. The three most upregulated KEGG pathways are *Ribosome, Steroid biosynthesis* and *Terpenoid backbone biosynthesis* (Fig. 4C). The three most downregulated KEGG pathways are *Axon guidance, Wnt signalling pathway* and *Synaptic vesicle cycle* (Fig. 4D).

**Fig. 4:**
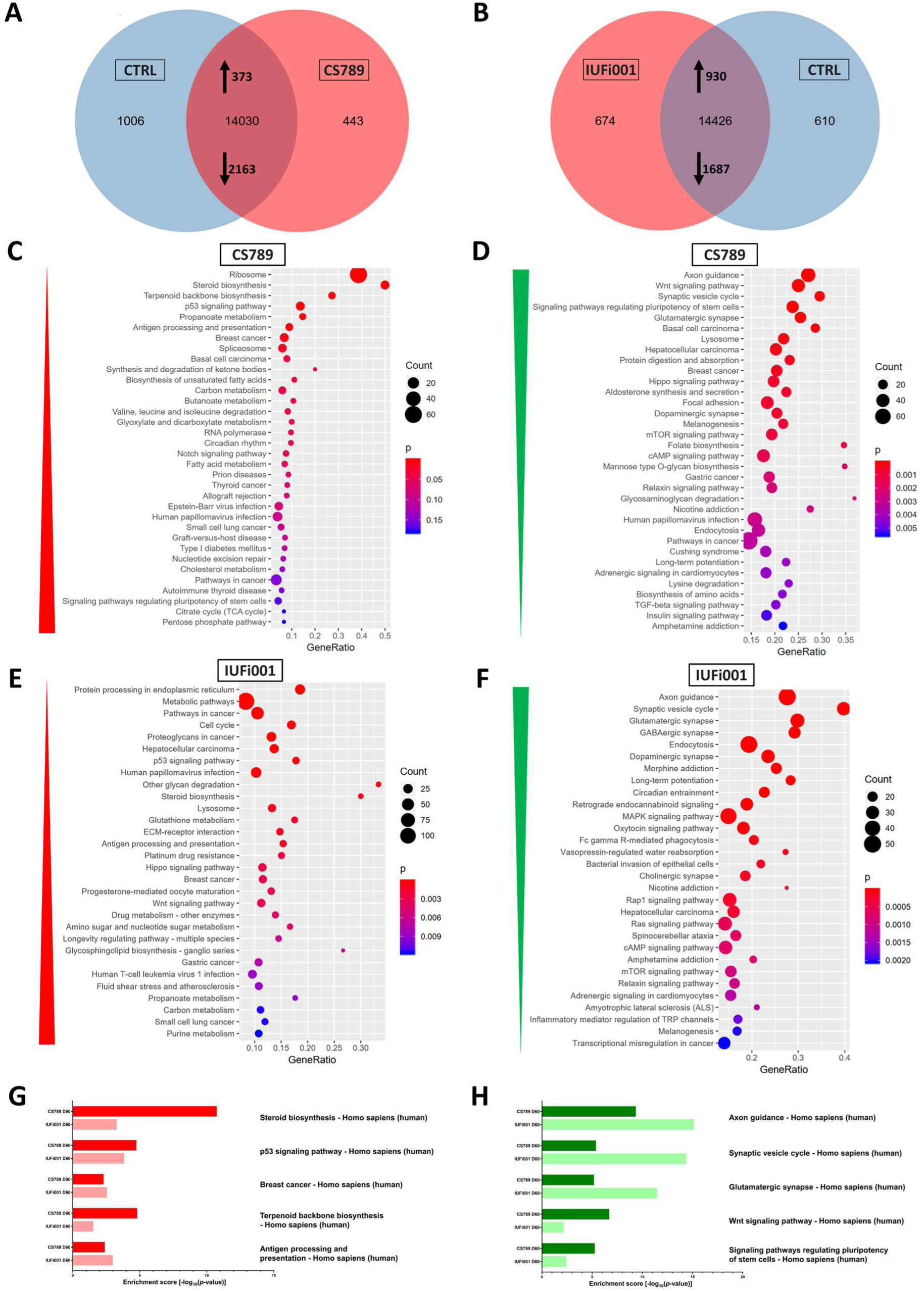
Global transcriptome and associated pathway analysis of control and CS organoids at day 60. (**A**) Venn diagram showing genes expressed only in CS789 organoids (443), in CTRL (B4) organoids (1006) and common to both (14030) (detection p value < 0.05). (**B**) Venn diagram showing genes expressed only in IUFi001 organoids (674), in CTRL (B4) organoids (610) and common to both (14426) (detection p value < 0.05). (**C,D**) Dot plots showing the Top 30 differentially regulated KEGG pathways (C) in the 373 significantly upregulated DEGs in day 60 CS789 organoids in comparison to CTRL (B4) (D) and in the 2163 significantly downregulated DEGs in day 60 CS789 organoids in comparison to CTRL (B4). (**E,F**) Dot plots showing the Top 30 differentially regulated KEGG pathways (E) in the 930 significantly upregulated DEGs in day 60 IUFi001 organoids in comparison to CTRL (B4) (F) and in the 1687 significantly downregulated DEGs in day 60 IUFi001 organoids in comparison to CTRL (B4). (**G**) Bar chart of the differentially upregulated KEGG pathways (Top 5 ranked) common between day 60 CS789 and IUFi001 organoids in comparison to CTRL (B4) organoids. (**H**) Bar chart of the differentially downregulated KEGG pathways (Top 5 ranked) common between day 60 CS789 and IUFi001 organoids in comparison to CTRL (B4) organoids.

Next, the transcriptome of the IUFi001 COs was compared to control COs. We identified 674 genes as exclusively expressed in IUFi001 COs, 610 genes exclusively expressed in control COs, and 14426 genes expressed in both. Of these 14426 commonly expressed genes, 930 genes are upregulated, and 1687 downregulated in IUFi001 organoids in comparison to control (Fig. 4B). The three most upregulated KEGG pathways are *Protein processing in endoplasmatic reticulum, Metabolic pathways* and *Pathways in cancer* (Fig. 4E). The five most downregulated KEGG pathways are *Axon guidance, Synaptic vesicle cycle* and *Glutamatergic synapse* (Fig. 4F).

We again compared the 50 highest up- and downregulated KEGG pathways in both patient-derived COs in comparison to control. The five most upregulated KEGG pathways common in CS789 and IUFi001 COs are *Steroid Biosynthesis, p53 signalling pathway, Breast cancer, Terpenoid backbone biosynthesis* and *Antigen processing and presentation.* (Fig. 4G). The five most downregulated KEGG pathways common in CS789 and IUFi001 COs are *Axon guidance, Synaptic vesicle cycle, Glutamatergic synapse, Wnt signalling pathway* and *Signalling pathways regulating pluripotency of stem cells*. (Fig. 4H). The complete list of pathways common in the 50 most severely dysregulated KEGG pathways can be found in Supplemental Table 2.

These results imply that while there are ample differences in gene expression between distinct individuals and severity grades of CS at later stages of BD, there is also severe common dysregulation, especially downregulation of genes important for BD and neuronal function. As with the neurospheres, this common dysregulation can already be detected in carriers of less severe forms of CS in early stages of BD.

### Non-redundant enrichment analysis reveals dysregulation of VEGFA-VEGFR2 signalling, brain development and intracellular transport in CSB-neurospheres

We performed analysis of the dysregulated Gene Ontologies (GOs) in our CS NS in comparison to control (Supplementary Figure S7). A variety of often related GOs were unveiled. To generate a clearer picture of the dysregulation in our CS neurospheres, the more extensive, upregulated gene-sets (CS789 2302 genes; IUFi001 1540 genes) were subjected to Metascape-based analysis, which resulted in non-redundant enrichment clusters. An analysis of the downregulated gene-sets can be found in Supplementary Figure S9.

For the CS789 NS, the three most enriched clusters were *VEGFA-VEGFR2 signalling pathway*, *cell morphogenesis* and *head development* (Fig. 5A). For the IUFi001 NS, the three most enriched clusters were *Vesicle-mediated transport*, *intracellular protein transport* and *Protein processing in endoplasmatic reticulum* (Fig. 5B).

**Fig. 5:**
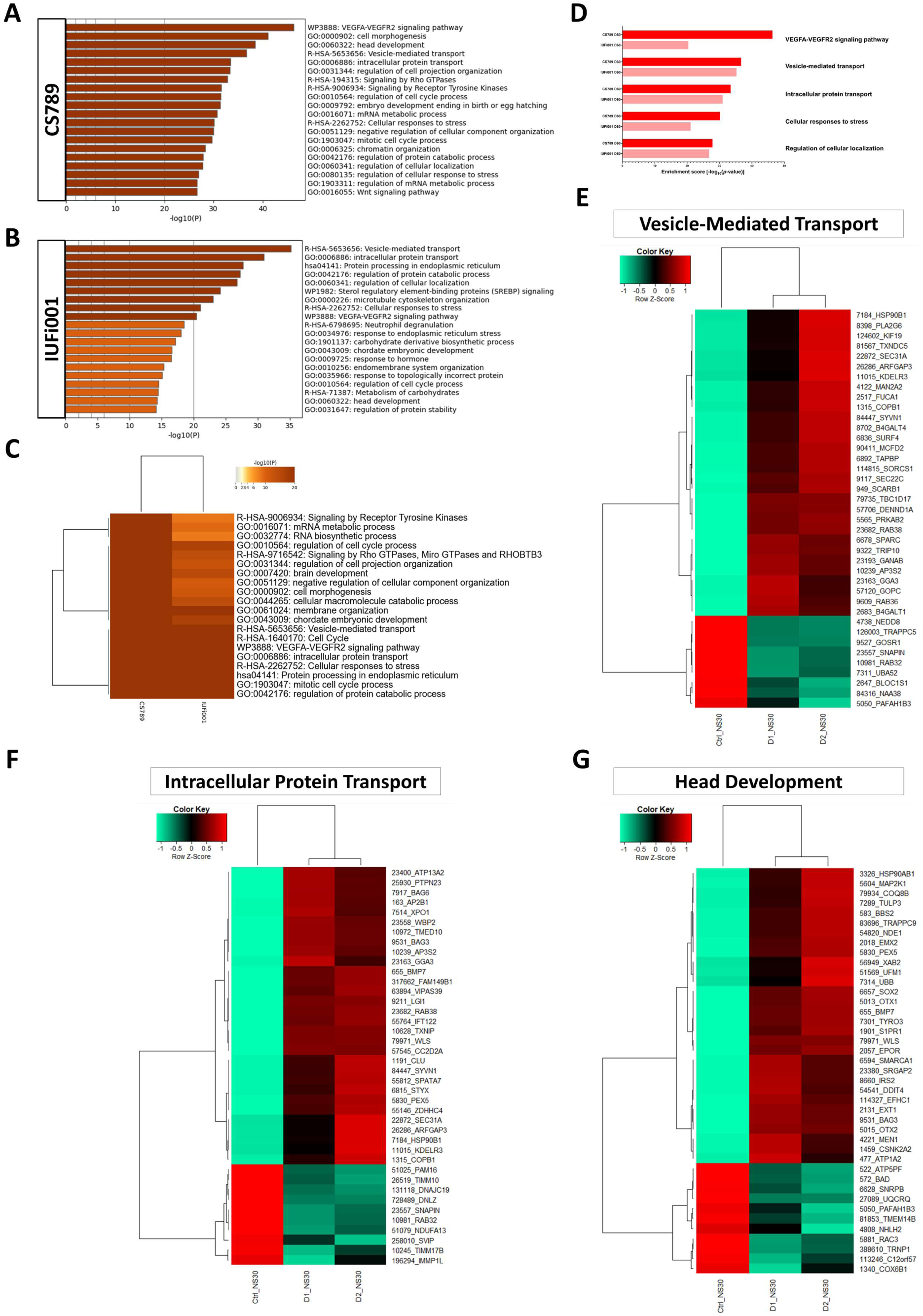
Comparative transcriptome and Gene Ontology analysis of upregulated DEGs in day 30 neurospheres. (**A**) Bar graph of the Top 20 non-redundant enrichment clusters attributable to the 2302 DEGs upregulated in day 30 CS789 neurospheres in comparison to CTRL (B4). (**B**) Bar graph of the Top 20 non-redundant enrichment clusters attributable to the 1540 DEGs upregulated in day 30 IUFi001 neurospheres. (**C**) Metascape-generated heatmap comparing upregulated gene-sets employed in A and B revealed i.a. GOs involved in brain development, intracellular transport and cell cycle. (**D**) Bar chart of the differentially upregulated enrichment clusters (Top 5 ranked) common between day 30 CS789 and IUFi001 neurospheres in comparison to CTRL (B4) neurospheres. (**E**) Pearsońs heatmap depicting the 40 most dysregulated genes involved in vesicle-mediated transport common between day 30 CS789 and IUFi001 neurospheres in comparison to CTRL (B4). (**F**) Pearsońs heatmap depicting the 40 most dysregulated genes involved in intracellular protein transport common between day 30 CS789 and IUFi001 neurospheres in comparison to CTRL (B4). (**G**) Pearsońs heatmap depicting the 40 most dysregulated genes involved in head development common between day 30 CS789 and IUFi001 neurospheres in comparison to CTRL (B4). (E-G) Pearsońs heatmaps depicting all dysregulated genes identified as common between CS789 and IUFi001 in these three GOs can be found in Supplementary Figure S11.

To pinpoint pathways of interest, we compared both gene-sets employing Metascape-based analysis. The highest consensus can be observed in *Vesicle-mediated transport*, *Cell cycle*, *VEGFA-VEGFR2 signalling pathway*, *intracellular protein transport* and *Cellular responses to stress* (Fig. 5C). Manual comparison of the singular analyses in Fig. 5A and Fig. 5B revealed identical enrichment clusters as the Metascape-based comparison (Fig. 5D).

Next, we chose the Reactome set *Vesicle-mediated transport* and the GOs *intracellular protein transport* and *head development*, compared the corresponding gene lists of both patient-derived NS and extracted the genes similarly regulated in comparison to control. This resulted in 165 genes for *intracellular protein transport*, 139 genes for *Vesicle-mediated transport* and 82 genes for *head development*. The resulting gene sets were compared utilizing Pearson’s heatmap analysis (Supplementary Figure S11). These gene sets were then reduced to the 40 highest dysregulated genes common between both patient-derived NS (Fig. 5E-G).

These results imply that at the developmental stage of NPCs there is severe dysregulation pertaining to distinct intracellular transport mechanisms, coordination of cell cycle and protein metabolism common between distinct grades of severity in CS patients.

### Non-redundant enrichment analysis reveals dysregulation of neuron projection development, brain development and distinct synaptic pathways in CSB-organoids

Furthermore, we performed analysis of the dysregulated GOs in our CS COs in comparison to control (Supplementary Figure S8). Again, a variety of often related GOs were unveiled. To produce a clearer picture of the dysregulation, the more extensive, downregulated gene-sets (CS789 2163 genes; IUFi001 1687 genes) were subjected to Metascape-based analysis, which results in non-redundant enrichment clusters. An analysis of the upregulated gene-sets can be found in Supplementary Figure S10.

For the CS789 COs, the three highest rated enrichment clusters were *neuron projection development, brain Development* and *regulation of neuron projection development* (Fig. 6A). For the IUFi001 COs, the three highest rated enrichment clusters were *neuron projection development, regulation of trans-synaptic signalling* and *brain development* (Fig. 6B). To pinpoint pathways of interest, we compared both gene sets employing Metascape-based analysis. The highest consensus can be observed in *brain development*, *Neuronal System*, *cell junction organization, regulation of trans-synaptic signalling* and *regulation of synapse structure or activity* (Fig. 6C). Manual comparison of the singular analyses in Fig. 6A and Fig. 6B revealed identical enrichment clusters as the Metascape-based comparison (Fig. 6D).

**Fig. 6:**
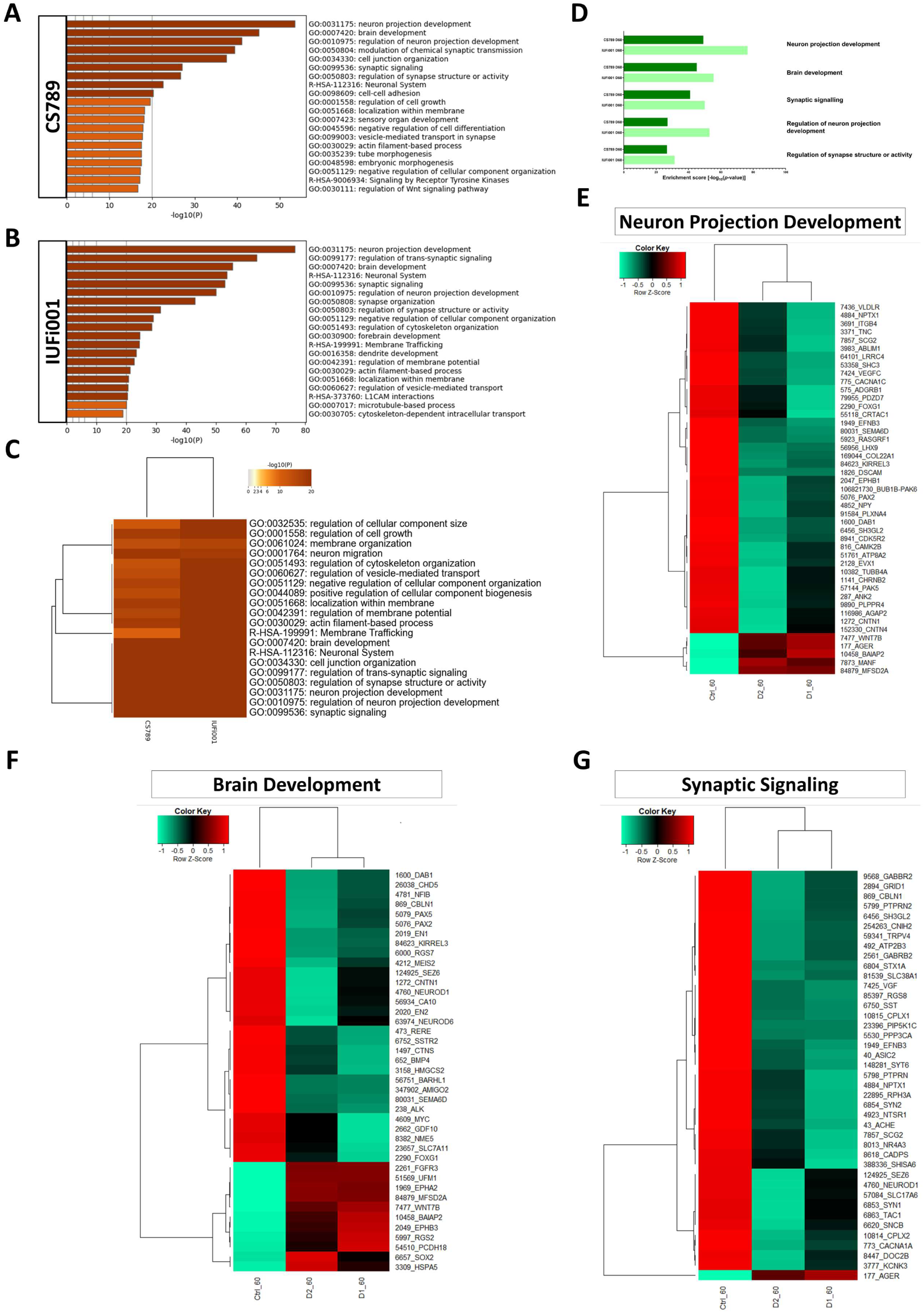
Comparative transcriptome and Gene Ontology analysis of downregulated DEGs in day 60 organoids. (**A**) Bar graph of the Top 20 non-redundant enrichment clusters attributable to the 2163 DEGs downregulated in day 60 CS789 (D1) organoids in comparison to CTRL (B4). (**B**) Bar graph of the Top 20 non-redundant enrichment clusters attributable to the 1687 DEGs downregulated in day 60 IUFi001 organoids. (**C**) Metascape-generated heatmap comparing downregulated gene-sets employed in A and B revealed i.a. GOs involved in brain development, cell junction organization, neuron projection development and synaptic signalling. (**D**) Bar chart of the differentially downregulated enrichment clusters (Top 5 ranked) common between day 30 CS789 and IUFi001 neurospheres in comparison to CTRL (B4) neurospheres. (**E**) Pearsońs heatmap depicting the 45 most dysregulated genes involved in neuron projection development common between day 60 CS789 and IUFi001 organoids in comparison to CTRL (B4). (**F**) Pearsońs heatmap depicting the 40 most dysregulated genes involved in brain development common between day 60 CS789 and IUFi001 organoids in comparison to CTRL (B4). (**G**) Pearsońs heatmap depicting the 40 most dysregulated genes involved in synaptic signalling common between day 60 CS789 and IUFi001 organoids in comparison to CTRL (B4). (**E-G**) Pearsońs heatmaps depicting all dysregulated genes identified as common between CS789 and IUFi001 in these three GOs can be found in Supplementary Figure S12.

Subsequently, we chose the GOs -*neuron projection development, synaptic signalling,* and *brain development*, compared the corresponding gene-sets of both patient-derived COs, and extracted the genes dysregulated in both patient-derived cell lines in comparison to control. This resulted in 165 genes for *neuron projection development*, 131 genes for *synaptic signalling* and 117 genes for *brain development*. The resulting gene-sets were compared utilizing Pearson’s heatmap analysis (Supplementary Figure S12). These gene-sets were then reduced to the 40 highest regulated genes common between both patient-derived COs (Fig. 6 E-G).

These results indicate that at the developmental stage of early BD there is severe dysregulation pertaining to neuron projection development, synaptic function and maintenance and even overall BD.

### Non-redundant enrichment analysis of genes commonly regulated at both timepoints reveals steroid biosynthesis as most severely affected

We were interested in genes and associated pathways commonly dysregulated at both developmental timepoints. We pooled the RNA-Seq data of both patient-derived samples at the NS and CO stage and compared them to their respective control via Fisheŕs exact test to determine the significantly regulated genes at both timepoints. Next, we compared the resulting gene-sets to find genes commonly dysregulated between both timepoints.

The list of commonly dysregulated genes was then subjected to Metascape-based analysis (Fig. 7A). The highest commonly dysregulated enrichment cluster is the KEGG pathway *Steroid Biosynthesis* and partially overlapping with it, in sixth and eighth place, the Reactome set *Activation of gene expression by SREBF (SREBP)* and the GO *fatty acid metabolism*. Other than that, there are three enrichment clusters we examined before, namely the GOs *synaptic signalling* and *brain development*, as well as the Reactome set *COPII-mediated vesicle transport*.

**Fig. 7:**
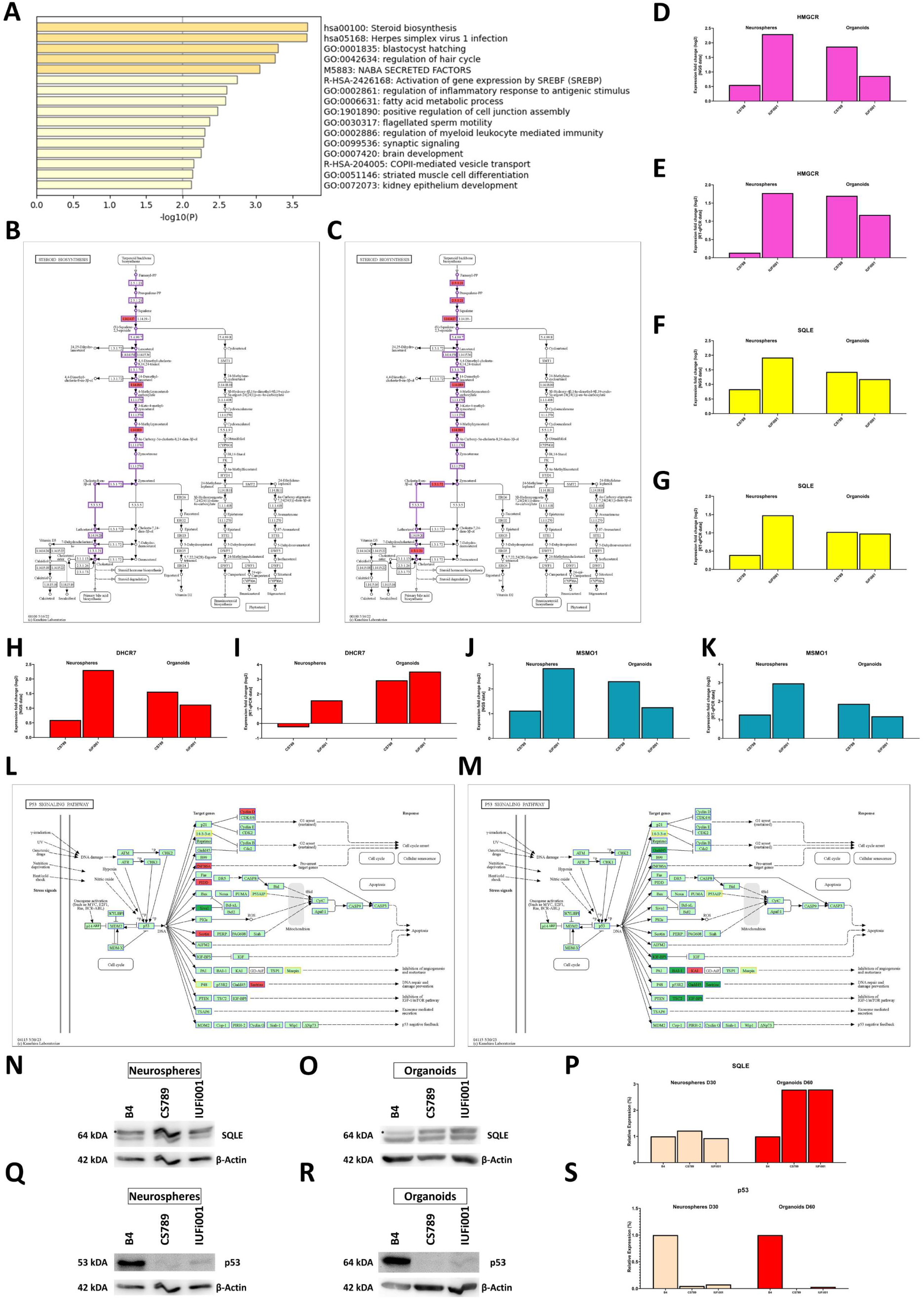
Metascape-based analysis of common genes dysregulated in day 30 CS neurospheres and day CS 60 organoids. (**A**) Bar graph of the 16 enrichment clusters attributable to the 181 DEGs regulated in both day 30 CS NS and day 60 CS COs in comparison to CTRL (B4). (**B, C)**) Schematic of the pathway *Steroid Biosynthesis* with the genes upregulated in CSB-deficient day 30 NS (B) and day 60 COs (C) indicated in red. (**D, F, H, J**) Relative mRNA expression analysis of *HMGCR* (D), *SQLE* (F), *DHCR7* (H), and *MSMO1* (J) in CS789 and IUFi001 organoids compared to CTRL (B4). (**E, G, I, K**) qRT-PCR analysis of *HMGCR* (E), *SQLE* (G), *DHCR7* (I), and *MSMO1* (K) mRNA expression in CS789 and IUFi001 organoids relative to CTRL (B4). (**L, M**) Schematic of p53 signalling pathway with genes upregulated in CSB-deficient day 30 NS (L) and day 60 COs (M) indicated in red and genes downregulated indicated in green. Gene which are not expressed are indicated in yellow. (**N, O**) Western blot analysis for SQLE in day 30 NS (N) and day 60 COs (O). (**P**) Quantification of SQLE western blot analysis at the day 30 NS and day 60 CO stage in CTRL (B4), CS789, and IUFi001. SQLE expression of CS789 and IUFi001 is compared to SQLE expression in CTRL (B4) of the respective timepoint. (**Q, R**) Western blot analysis for p53 in day 30 NS (Q) and day 60 COs (R). (**S**) Quantification of p53 western blot analysis at the day 30 NS and day 60 CO stage in CTRL (B4), CS789, and IUFi001. SQLE expression of CS789 and IUFi001 is compared to p53 expression in CTRL (B4) of the respective timepoint.

Next, we investigated which of the commonly dysregulated genes are most severely regulated in both patient-derived lines at both timepoint in comparison to control. We found that the genes MAGE Family member A4 (*MAGEA4*), Transmembrane Protein 132C (*TMEM132C*), Zinc Finger Protein 558 (*ZNF558*) and Tripartite Motif Containing 4 (*TRIM4*) are the most severely and consistently regulated genes in all patient-derived samples (Supplementary Figure S13 A, C, E, G).

After identifying these four genes, we ascertained our findings via RT-qPCR (Supplementary Figure S13 B, D, F, G). We further investigated these four genes in another sets of day 60 COs, produced employing an altered media composition. In these COs, we detected regulation of *MAGEA4*, *TMEM132C*, *ZNF558* and *TRIM4* akin to our initial dataset (Supplementary Figure S13 H-K). We cultivated a small number of COs from the initial batch investigated in this work until day 120. RTqPCR of these samples also revealed regulation of these four genes (Supplementary Figure S13 L-O). Following up on our initial Metascape-based analysis, we identified regulated genes involved in cholesterol biosynthesis (CB) in the KEGG pathway *Steroid Biosynthesis* at both timepoints. As CB requires the metabolite Farnesyl pyrophosphate, we simultaneously investigated the Mevalonate pathway branch of *Terpenoid Backbone Synthesis* at both timepoints. While we found no significantly regulated genes in the NS, analysis of the CO datasets revealed regulation of Hydroxymethylglutaryl-CoA Synthase (*HMGCS*), Hydroxymethylglutaryl-CoA Reductase (*HMGCR*), one of the rate-limiting enzymes of CB, as well as Isopentenyl-Diphosphate Delta-Isomerase (*IDI*) and Farnesyl Diphosphate Synthase (*FDPS*) (Supplementary Figure S14). Analysis of the CB branch of the KEGG pathway *Steroid Biosynthesis* indicated the second rate-limiting enzyme of CB Squalene Epoxidase (*SQLE*) and Methylsterol Mono-oxygenase (*MSMO1*) as regulated in both NS and COs. Furthermore, in the COs, the genes Farnesyl-Diphosphate Farnesyltransferase 1 (*FDFT1*), 7-Dehydrocholesterol Reductase (*DHCR7*) and 24-Dehydrocholesterol Reductase (*DHCR24*) were found to be differentially regulated (Fig. 7 B, C).

We validated these findings for the genes *HMGCR*, *SQLE*, *DHCR7* and *MSMO1* via RT-qPCR (Fig. 7 D-K). We further analysed, whether the regulation of *SQLE* translated to an increase in protein levels via western blot. We found no difference in the NS, but a nearly threefold increase of SQLE in the COs (Fig. 7 N-P). The full Western Blots can be found in (Supplementary Figure S15).

Next, we analysed a pathway known to be dysregulated in CSB-deficient cells, *p53 signalling pathway*. In the NS, we found upregulation of Zinc Finger Protein 385A (*ZNF385A*), P53-Induced Death Domain Protein 1 (*PIDD1*), Shisa Family Member 5 (*SHISA5*), Cyclin D1 (*CCND1*), Sestrin 2 (*SESN2*) and Sestrin 3 (*SESN3*), as well as downregulation of SIVA1 Apoptosis Inducing Factor (*SIVA1*). In the COs, we found downregulation of Growth Arrest and DNA Damage Inducible Gamma (*GADD45G*), Insulin Like Growth Factor Binding Protein 3 (*IGFBP3*), Adhesion G Protein-Coupled Receptor B1 (*ADGRB1*), *SESN2* and TSC Complex Subunit 2 (*TSC2*) as well as upregulation of CD82 Molecule (*CD82*) (Fig. 7 L, M). We further analysed the quantity of p53 protein at both timepoints. Contrary to our expectations, we detected a decrease in p53 protein levels at both timepoints (Fig. 7 Q-S). The full Western Blots can be found in Supplementary Figure S15.

These results indicate upregulation of CB during early cerebral development of CS patients. While we could identify transcriptional upregulation of genes involved in CB in both day 30 NS and day 60 COs, this upregulation could only be confirmed on the translational level in the COs. Our results also confirm the dysregulation of the p53 signalling pathway in iPSC-derived NPCs and neurons during early human BD in CS patients. Interestingly, this disruption of p53 signalling pathway manifests in altered expression of different genes between day 30 NS and day 60 COs. Unexpectedly, we detected a decrease of p53 protein levels at both timepoints.

## Discussion

The major diagnostic criteria of CS are mostly symptoms of the central nervous system, e.g., progressive microcephaly, neurologic dysfunction, and cerebellar atrophy. However, the mechanisms underlying these manifestations are still largely unknown, due to the versatility of the causative protein and limitations of the available experimental models. To overcome this deficiency, we produced the first successful differentiation of CSB-deficient iPSC lines into cerebral organoids (COs). We employed two distinct, patient-derived, CSB-deficient iPSC lines, one generated from a donor with the most severe form of CS, cerebro-oculo-fascio-skeletal syndrome (COFS), and one generated from a donor with classical CS Type I and differentiated them into neural progenitor cells (NPCs), in the form of neurospheres (NS), and COs. To elucidate the common dysregulation underlying aberrations in brain development (BD) found in different types of CS, we then subjected mRNA of both developmental timepoints, day 30 NS and day 60 COs, to NGS and the resulting datasets to extensive bioinformatic analysis. The transcriptome of day 30 NS is estimated to correspond to 8-9 week post-conception fetal brains and the transcriptome of day 60 COs to 13-21 week post-conception fetal brains.^32,49^ COs produced with this protocol have been shown to consist mostly of cells with forebrain identity, however sub-populations of cells with midbrain and hindbrain identity are to be expected.^32,50^

Transcriptome-based analysis of the differentially regulated KEGG pathways in the NS revealed several common pathways. The upregulated pathway *Protein processing in endoplasmatic reticulum* and the downregulated pathway *Ribosome* could indicate changes in the translational priming of NPCs. Ribosome biogenesis has been shown to be impaired in CSA- and CSB-deficient cells before, due to a disturbance of RNA polymerase I transcription and processing of pre-rRNA.^51,52^ A decrease in ribosome-related genes would be expected to result in a decrease in protein processing. This might indicate a compensatory mechanism to increase global protein production. However, it could also indicate dysregulation of global protein synthesis, which has been implicated in disease progression of schizophrenia patients.^53^

Furthermore, our analysis indicated *VEGFA-VEGFR2 signalling pathway* as modulated in our patient-derived cell lines. While known for its function in angiogenesis, VEGFA also has direct impact on NPCs and neurons. VEGFA has been shown to increase proliferation of primary embryonic cortical NPCs.^54,55^ VEGFA has also been shown to decrease apoptosis in adult NPCs, and to promote neuronal survival.^56–60^ Additionally, VEGFA is surmised to be involved in dendrite outgrowth and axon guidance, two processes we found regulated in day 60 COs.^55,57^ Overall, this increase in VEGFA-VEGFR2 signalling might present a compensatory mechanism to increase proliferation, which has been shown to be reduced in CSB-deficient NPCs^61^ or a protective response to increased ambient stress, which can be observed in the form of *Cellular responses to stress*.

We also provide an in-depth analysis of several regulated enrichment clusters in our patient-derived NS via heatmap analysis, namely *Vesicle-mediated transport*, *intracellular protein transport* and *head development*. Vesicular transport is an essential process by which membrane-bound vesicles are released from a donor compartment and transported to a specific cellular location. Disturbances of *Vesicle-mediated transport* cause a multitude of disorders, several of which share symptoms with CS.^62^ The following genes identified as regulated in our dataset have been associated with diseases caused by alterations in *Vesicle-mediated transport*: *LMAN1, SEC23A, SEC23B, SEC24B, SEC31A, GAK, VPS35L, DNM2, SBF1, BLOC1S1, COG1, COG4, SCYL1, MTM1, AP1S2, DENND5A, TRAPPC*9.^63^ The dysregulation of a subpathway, namely *COPII-mediated vesicle transport*, is also observable between both investigated timepoints.

We were further interested in the GO *head development*. The observed upregulation of BD it encompasses could indicate a premature entry of NPCs into neurogenesis, a hallmark of microcephaly. One reason for this premature shift could be the increase in ambient cell stress observed in our analysis, as DNA damage can promote accelerated neurogenesis.^32,64–66^ Interestingly, CS patients, except for COFS patients, do not present microcephaly in prenatal development. Still, both of our patient-derived cell lines display similar regulation of BD at the NPC stage, indicating potential issues with developmental timing.

The highest upregulated KEGG pathway in day 60 COs, *Steroid Biosynthesis*, shows common dysregulation over both timepoints. *Steroid Biosynthesis,* as well as the commonly dysregulated Reactome set *Activation of gene expression by SREBF (SREBP)*, both encompass cholesterol biosynthesis (CB). Cholesterol is produced by neurons at early stages of BD. Cholesterol is necessary for the control of membrane fluidity and lipid raft structure and influences e.g., neuronal receptor activity.^67–69^ Correctly orchestrated CB is essential for BD and directly influences pathways dysregulated in our CSB-deficient lines, as cholesterol is involved in synaptogenesis, stability and recycling of synaptic neurotransmitters and neurite outgrowth.^68,70–73^ Cholesterol depletion, as well as accumulation can lead to neuronal dysfunction and ultimately to neurodegeneration.^72,74^ In a mouse model with conditional knock-out of CB in radial glia cells, the affected pups exhibited progressive loss of cortical neurons, as well as postnatal proliferation and migration deficits of cerebellar granule precursors.^75^ Overall, diseases of cholesterol metabolism have a high overlap of symptoms with CS, with many manifesting with i.a. neurological symptoms like psycho-motor retardation, developmental delay, and microcephaly.^76^

We verified this upregulation of CB in the COs via RT-qPCR of the two rate-limiting enzymes *HMGCR* and *SQLE*, as well as *DHCR7* and *MSMO1*. On the protein level, we could detect a nearly threefold increase of SQLE protein, reinforcing the hypothesis that CB is increased in CSB-deficient young neurons. Interestingly, while we could also detect an increase in *SQLE* transcription in our NS, this did not translate into an increase in SQLE protein. However, NPCs react differently to cholesterol depletion than neurons. In a mouse model with a conditional *FDFT1* knock-out in VZ-NPCs, VEGF expression via a hypoxia-inducible factor-1 independent pathway was strongly upregulated. This increased angiogenesis and thus influx of external cholesterol to the NPCs.^77^ Fittingly, we also identified upregulation of *VEGFA-VEGFR2 signalling pathway* in our NS.

The *p53-signalling pathway*, reputed in its role as a tumour suppressor and its function in DNA damage response, cell cycle arrest, senescence, and apoptosis, is also involved in BD. Deletion, mutation, and inhibition of p53 can lead to female-biased neural tube defects.^78^ However, deletions of effectors of p53 can lead to various brain malformations.^79^ Due to the variety of potential outcomes, it is surmised, that p53 stabilization has a controlled, dose-and-time dependent effect on distinct cell types during development.^78^ In the context of CS, CSA and CSB have been shown to interact with p53 and regulate its degradation. Both proteins form a complex with p53 and MDM2, which facilitates ubiquitination and subsequent degradation of p53 in an MDM2-dependent manner.^80^ Due to decreased degradation, CSA- and CSB-deficient cells show elevated levels of p53.^81^ CSB also interacts directly with p53, competing with the essential factor E1A Binding Protein p300 (*EP300*) for its binding site. As CSB has higher affinity for p53, this negatively modulates the function of p53.^82–84^ In summary, regulation of *p53 Signalling pathway* was expected, due to a decrease in degradation of p53 and absence of a negative regulator of p53 activity. This might have implications for BD, as p53 is involved in proliferation and differentiation of NPCs^85^ and controlled dose- and time-dependent stabilization of p53 is necessary for correct development.

Surprisingly, instead of the expected increase, we detected a decrease in p53 protein levels at both timepoints. p53 protein level have been revealed to be high during early embryogenesis and to decrease over the course of development until it plateaus at a low level in terminally differentiated cells.^86–88^ This decrease of p53 protein levels we identified might be another manifestation of the dysregulation of BD we revealed in this work, further uncoupling development of our CS-deficient cells from normal cerebral development. Another possibility would be increased modification of p53, depleting the pool of unmodified p53 we examined, which would also directly impact BD.

The most significantly downregulated KEGG pathway in both patient-derived COs, *Axon Guidance*, has been shown to be dysregulated in human neuronal networks, as well as *Axonogenesis*, which shows in our analysis as part of *Neuron projection development*.^26^ In human neural cell lines, ablation of CSB expression affected neuronal differentiation capabilities. Suppression of CSB led to a decrease in MAP2 expression, accompanied by reduced cell polarization. Together this led to a decrease in neuritogenesis.^15,89^ Alterations of neuritogenesis and axonal pathfinding may lead to pathological changes in neural circuitry.

The downregulated KEGG pathway *Synaptic Vesicle Cycle* has been demonstrated to be dysregulated, as well as other pathways involved in synapse formation, activity and maintenance, whose dysregulation we could observe with the regulated GOs *regulation of trans-synaptic signalling, regulation of synapse structure*, and *synaptic signalling.*^26^ Correct function of synapse pathways are necessary for neuronal function and signal transmission, which both have been shown to be altered in CSB-deficient neural networks.^26^ In CS patient cerebella, downregulation of genes involved in synaptic exocytosis has been identified, which we also identified in our COs.^15^ Our data also indicates downregulation of synaptic endocytosis in our patient-derived COs. This alteration of synapse formation, activity and maintenance and the resulting changes likely contribute to the neurodevelopmental defects in CS patients.

We further provide an analysis of several regulated enrichment clusters in our patient-derived COs via heatmap analysis, namely the GOs *neuron projection development, synaptic signalling,* and *brain development*.

*Brain development*, which was upregulated in NS, was found to be downregulated in day 60 COs. This was expected, as CSB-deficient cells have been shown to exhibit downregulation of thousands of genes, including brain-related genes, due to dysregulation of RNA Polymerase II activity.^14,15^ We found the *Brain development*-related genes *EPHA7, EPHB3, AFF2, NPY, WNT7B, TSPAN2, SRGAP2, UFM1, FUT10, POTEE, TRH* and *TPGS1* to be mostly upregulated in NS and downregulated in COs at both investigated timepoints.

However, the most significantly dysregulated genes are not associated with *brain development*. We found the genes *MAGEA4*, *TMEM132C*, *ZNF558* and *TRIM4* to be most severely regulated in all patient-derived samples. Other members of the MAGE-A family have been shown to potentially be involved in early BD.^90^ *TMEM132C* has been revealed to be expressed in early NPCs, the developing cochlea and is differentially regulated in dorsal forebrain and midbrain during murine development.^91^ *TMEM132C* has also been shown to be differentially regulated in NPCs of another disease with congenital cataracts and intellectual disability, Lowe syndrome.^92^ Krüppel-associated box (KRAB) Zinc

Finger Protein *ZNF558*, has a role in mitochondrial maintenance and is thought to influence timing during early human BD.^93^ *TRIM4* is a gene mainly known for its role in ubiquitination of a variety of targets.^94–96^ A genome wide DNA methylation study found that upregulation of *TRIM4* might be associated with neural tube defects.^97^

So, while these four genes are not yet conclusively linked to BD, one can assume they might be associated. Interestingly, this dysregulation of all four genes seems to be traceable to later timepoints, indicating their involvement not only in early, but at least also mid-gestational BD.

## Conclusions

In conclusion, this study is the first to produce CSB-deficient patient-derived 3D organoids. Our data provides new insights into the transcriptional dysregulation associated with brain development in CS patients. We confirm known dysregulation of neuron projection- and synapse-related pathways and expand this set with the dysregulation of the metabolic pathway *Steroid Biosynthesis*, as well as dysregulation of lipid metabolism in NPCs and neurons in a 3D environment. Our data further provides substantiation to the theory, that CS is not only a neurodegenerative, but also a neurodevelopmental disorder, with characteristic dysregulation of brain development at distinct stages. We also identified highly differentially regulated genes not previously highlighted in CSB-deficient neuronal cultures. These genes seem to be dysregulated at several stages of brain development, indicating them as potential target candidates for intervention. As a future prospect, a deeper understanding of the underlying developmental dysregulation in the most devastating symptoms of CS can hopefully spur further research and development of therapeutic strategies.

## Acknowledgements

The authors are grateful to Kanehisa Laboratories for the permission to use their figures for Terpenoid Backbone Biosynthesis, Steroid Biosynthesis and p53 signalling pathway for our figures. The figures were taken from https://www.genome.jp/kegg/. The authors are also grateful to Marcel Tisch (https://twitter.com/MarcelTisch) for producing the icons for Embryoid Bodies, Organoids and Stem Cell Colony used in the graphical abstract and allocating them to public domain. The icons can be found at https://bioicons.com.

## Author Contributions

L.S. designed and performed experiments, analysed data, wrote and edited the manuscript. W.W. performed the bioinformatic analysis, data curation, provided figures and edited the manuscript. J.Ka. performed cell banking. A.R. provided a cell line. E.F. acquired funding and provided guidance. J.Kr. acquired funding and provided guidance. J.A. conceptualized the work, wrote and edited the manuscript, coordinated the study, acquired funding and supervised the study. All authors have read and agreed to the published version of the manuscript.

