## Supplementary Figures S1-S15 for "Cockayne syndrome patient iPSC-derived brain organoids and neurospheres show early transcriptional dysregulation of biological processes associated with brain development and metabolism"

**A**

| | B4 | CS789 | IUFi001 |
|---|---|---|---|
| **Cell Type** | Fetal Foreskin Fibroblast | Dermal Fibroblast | Dermal Fibroblast |
| **Provided By** | ISRM Düsseldorf | IUF - Leibniz Research Institute for Environmental Medicine | |
| **Donor Age** | Neonatal | 10 months | 3 years |
| **Sex** | ♂ | ♂ | ♀ |
| **Reprogramming** | Viral | Episomal | Episomal |
| ***ERCC6* Mutation** | None | Exon 10/R683X | Exon 5/K377X; Exon15/R857X |
| **CS Severity** | None | COFS | CS Type I |

**B**

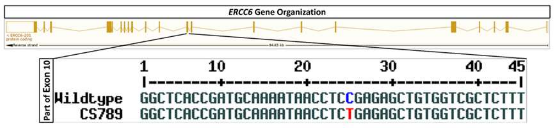


**C**

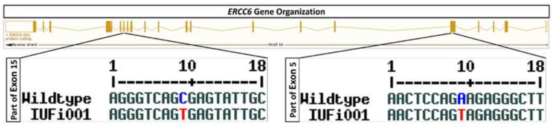


**D**

| | Cerebrooculofacioskeletal (COFS) syndrome | Cockayne Syndrome Type II | Cockayne Syndrome Type I | Cockayne Syndrome Type III | UV-sensitive Syndrome |
|---|---|---|---|---|---|
| **Onset** | Fetal | At birth | First 2 years | 3-4 years | Varying |
| **Diagnosed at** | Birth | Infancy | Childhood | Early teens | Childhood/Adulthood |
| **Life Expectancy** | Infancy | 5-6 Years | 16 years | 30 years | Normal? |
| **Suggestive Findings** | Athrogryphosis<br>Prenatal growth failure<br>Prenatal Microcephaly<br>Congenital Cataracts **or**<br>Congenital Microphthalmia | **Major Criteria:**<br>Postnatal growth failure; Progressive Microcephaly; Neurologic Dysfunction with developmental delay; White matter dysmyelination; Cerebellar Atrophy; Intracranial calcifications<br>**Minor Criteria:**<br>Cutaneous Photosensitivity; Demyelinating peripheral neuropathy; Pigmentary Retinopathy; Cataracts; Sensorineural Hearing loss; Enamel Hypoplasia; Tooth Anomalies; Cachectic Dwarfism | | | |
| **Diagnosis** | Multigene panel **or** Comprehensive genomic testing | | | | |

**Supplementary Figure S1: Used cell lines and clinical information about Cockayne Syndrome.** (**A**) General information about the cell lines used in this work. (**B**) Schematic depiction of the ERCC6 mutation found in the CS789 iPSC line. (**C**) Schematic depiction of the ERCC6 mutation found in the IUFi001 iPSC line. (**D**) Table showing clinical features and diagnostic criteria of different types of Cockayne Syndrome.

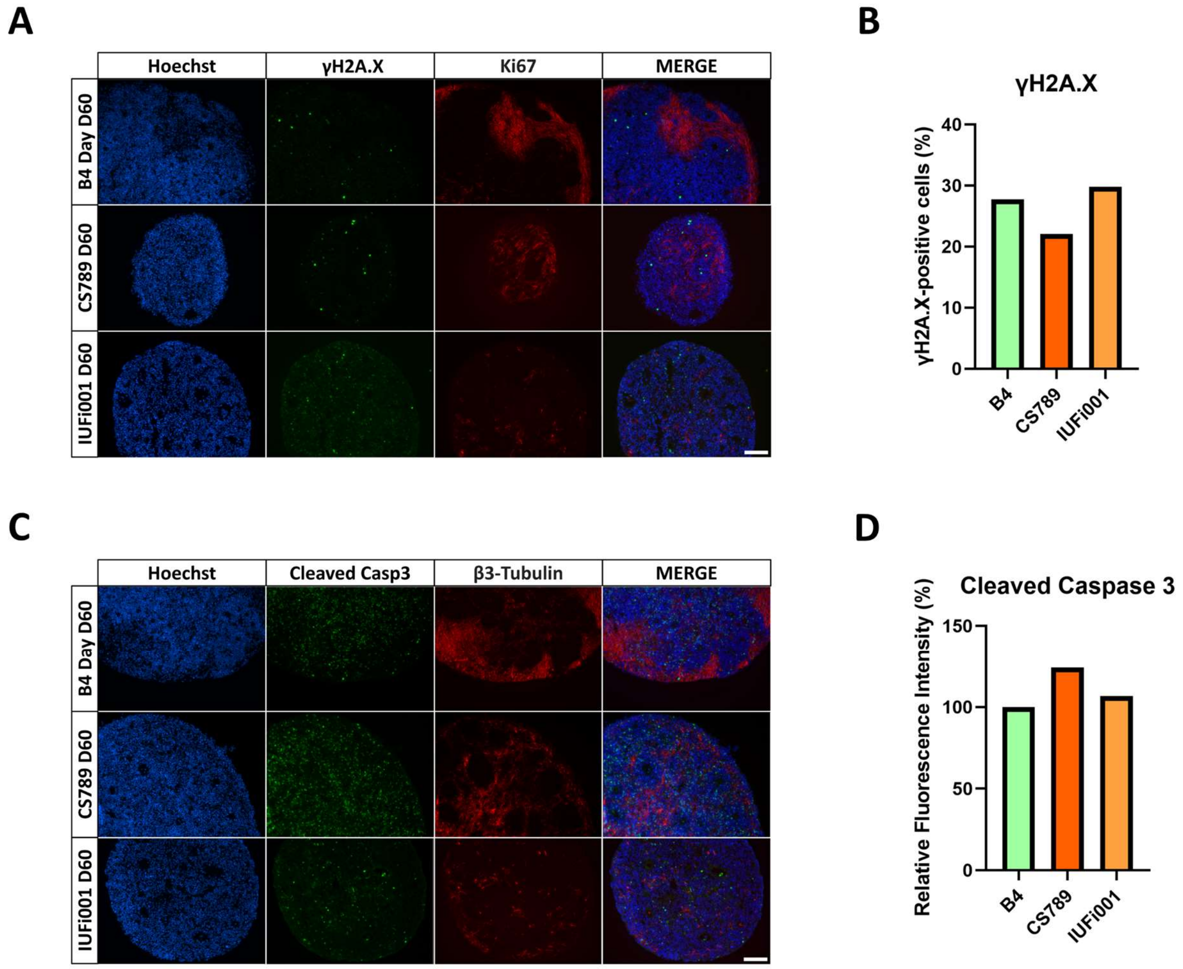


**Supplementary Figure S2: DNA damage-related marker γH2A.X and apoptosis marker C-Caspase 3 in day 30 neurospheres.** (**A**) Representative immunocytochemistry images of the distribution of cells expressing γH2A.X and β3-Tubulin. 200x magnification, scale bar 100 µm. (**B**) Quantification of the γH2A.X-positive, Hoechst-positive cells in CTRL (B4), CS789 and IUFi001 neurospheres. (**C**) Representative immunocytochemistry images of the distribution of cells expressing cleaved caspase 3 and β3-Tubulin. 200x magnification, scale bar 100 µm. (**D**) Quantification of the relative fluorescence intensity in CTRL (B4), CS789 and IUFi001 neurospheres. (**B,D**) At least ten random fields from three different neurospheres of every cell line were analysed.

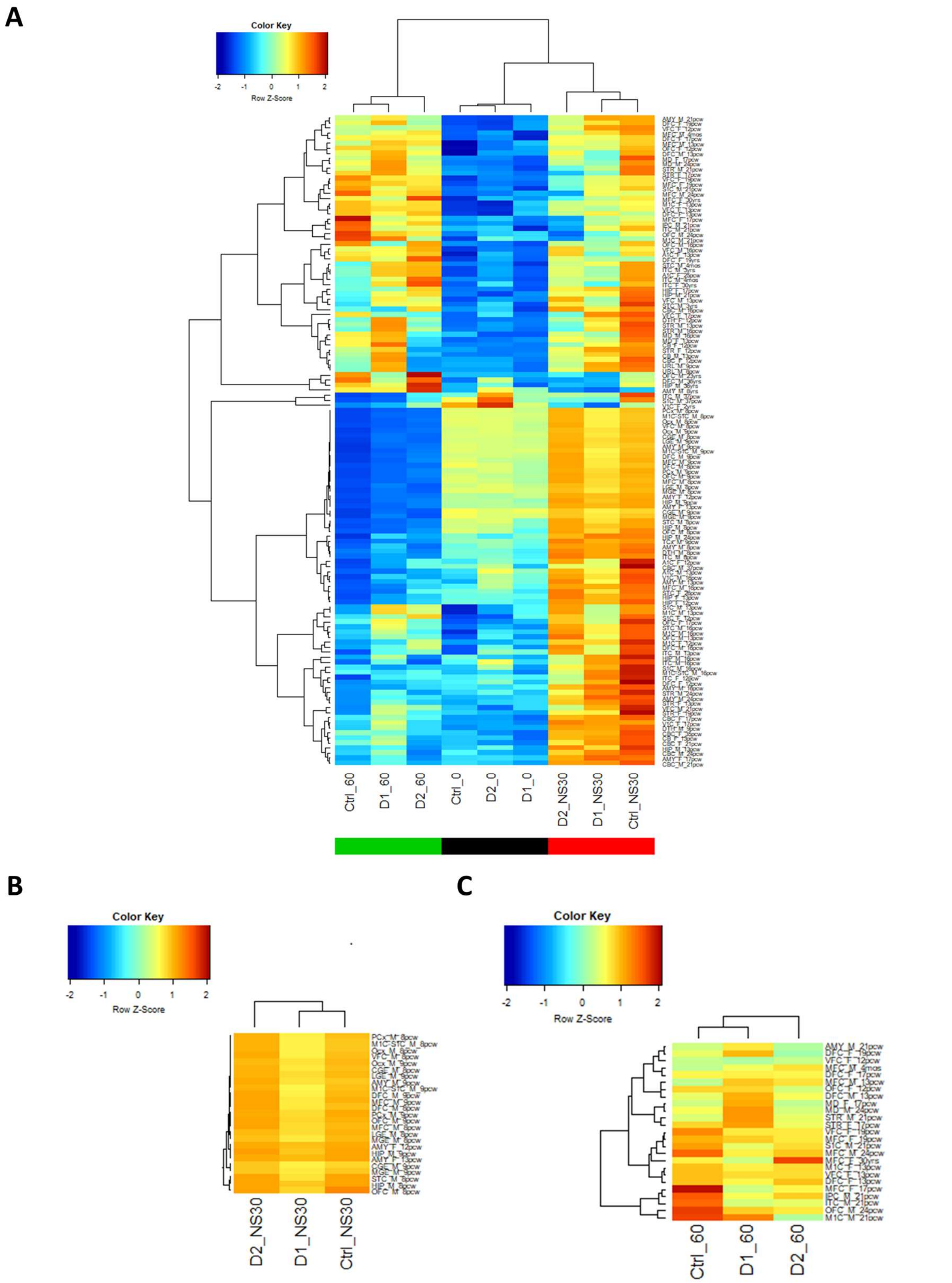


**Supplementary Figure S3: Determination of developmental age of day 30 neurospheres and day 60 cortical organoids.** (**A**) Heat map of enrichment scores of transcriptomic data from control and CS iPSCs, neurospheres and organoids in comparison with the Allen Brain Atlas (ABA) human developmental Brain NGS data (https://www.brainspan.org/). (**B**) Region of highest consensus for the day 30 neurospheres extracted from A. (**C**) Region of highest consensus for the day 60 organoids extracted from A.

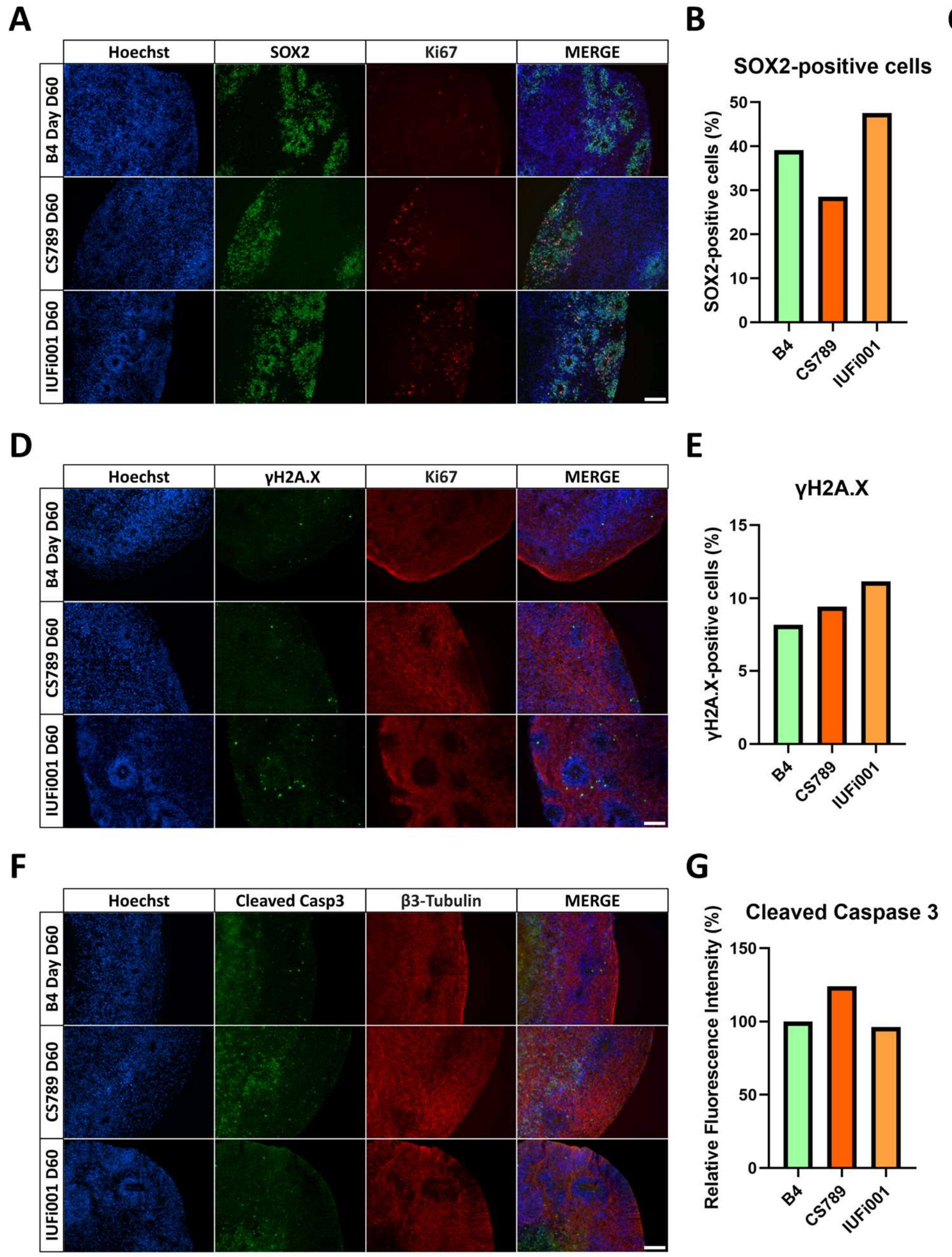


**Supplementary Figure S4: Characterization of the cellular composition and DNA damage response of day 60 organoids.** (**A**) Representative immunocytochemistry images of the distribution of cells expressing SOX2 and Ki67. 200x magnification, scale bar 100 µm. (**B**) Quantification of the SOX2-positive, Hoechst-positive cells in CTRL (B4), CS789 and IUFi001 neurospheres. (**C**) Quantification of the KI67-positive cells within the SOX2-positive cell population in CTRL (B4), CS789 and IUFi001 organoids. (**C**) Representative immunocytochemistry images of the distribution of cells expressing γH2A.X and β3-Tubulin in CTRL (B4), CS789 and IUFi001 organoids. 200x magnification, scale bar 100 µm. (**D**) Quantification of the γH2A.X-positive, Hoechst-positive cells in CTRL (B4), CS789 and IUFi001 organoids. (**E**) Representative immunocytochemistry images of the distribution of cells expressing cleaved caspase 3 and β3-Tubulin in CTRL (B4), CS789 and IUFi001 organoids. 200x magnification, scale bar 100 µm. (**F**) Quantification of the relative fluorescence intensity in CTRL (B4), CS789 and IUFi001 organoids. (**B,C,E,G**) At least ten random fields from three different organoids of every cell line were analysed.

**A**

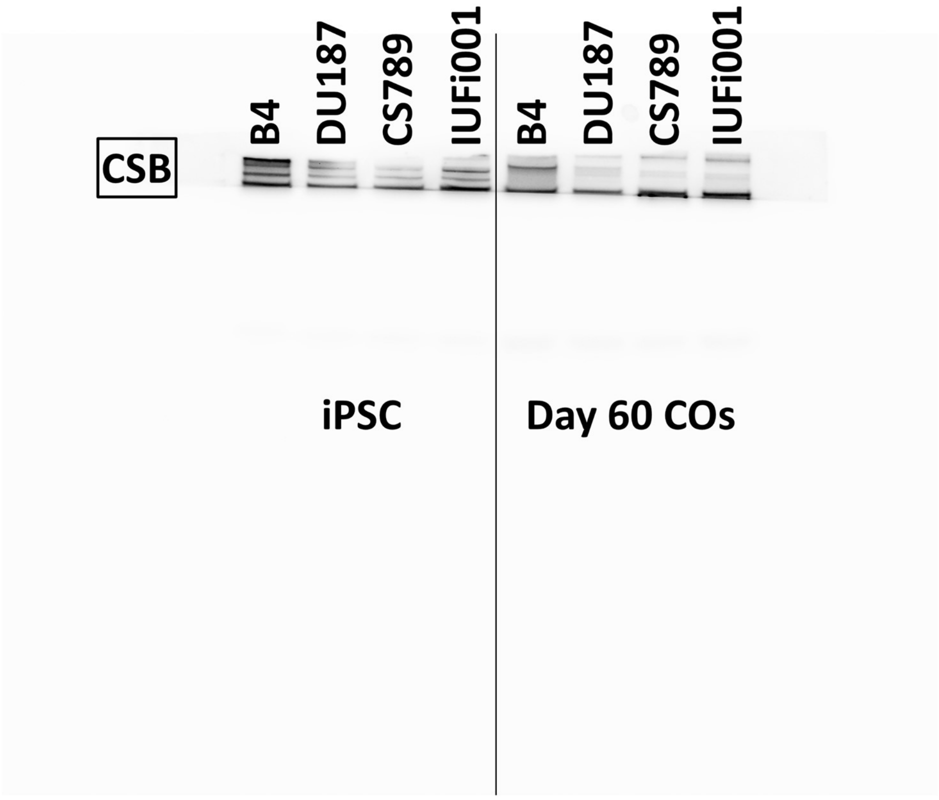


**B**

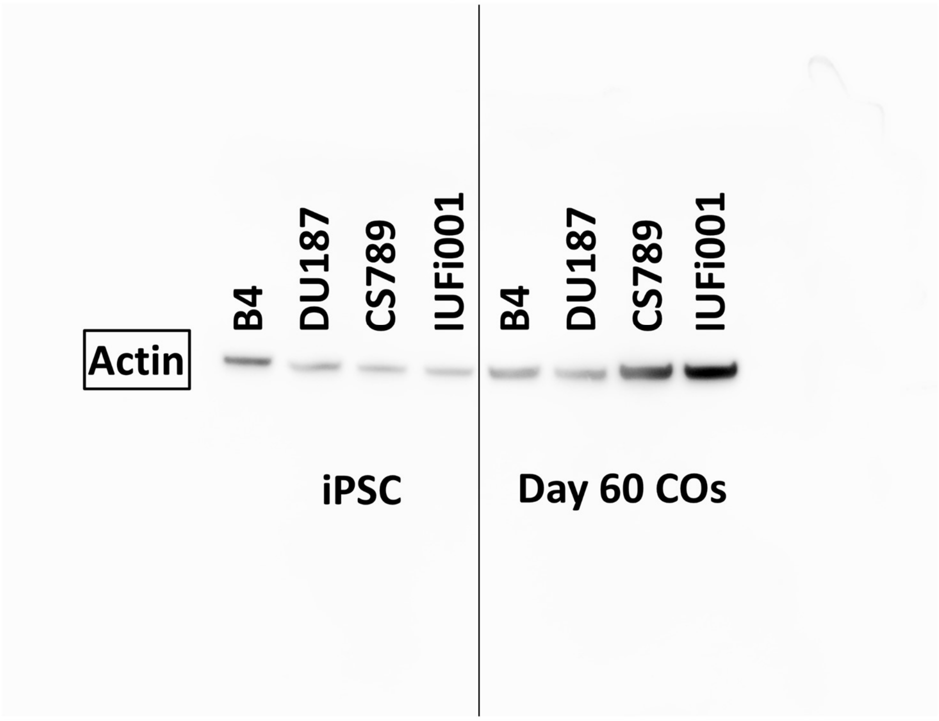


**Supplementary Figure S5: Full Western Blot for CSB protein and Actin protein in Day 0 iPSCs and Day 60 COs.** (**A**) Western Blot analysis for full-length CSB protein in B4, DU187, CS789 and IUFi001 iPSCs and COs. The cell line DU187 was not included in this article. (**B**) Western Blot for beta-Actin protein in B4, DU187, CS789 and IUFi001 iPSCs and COs. The cell line DU187 was not included in this article.

**A**

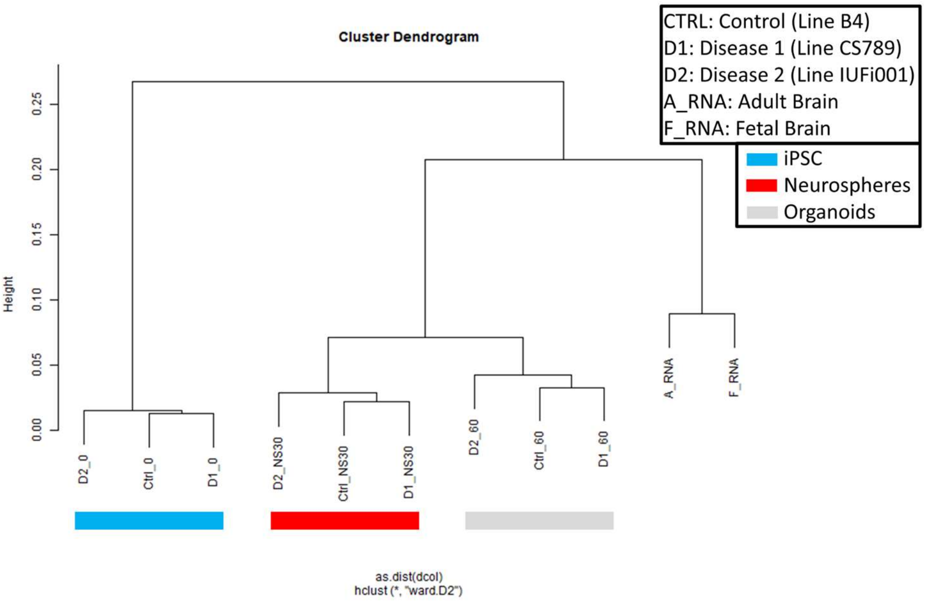


**B**

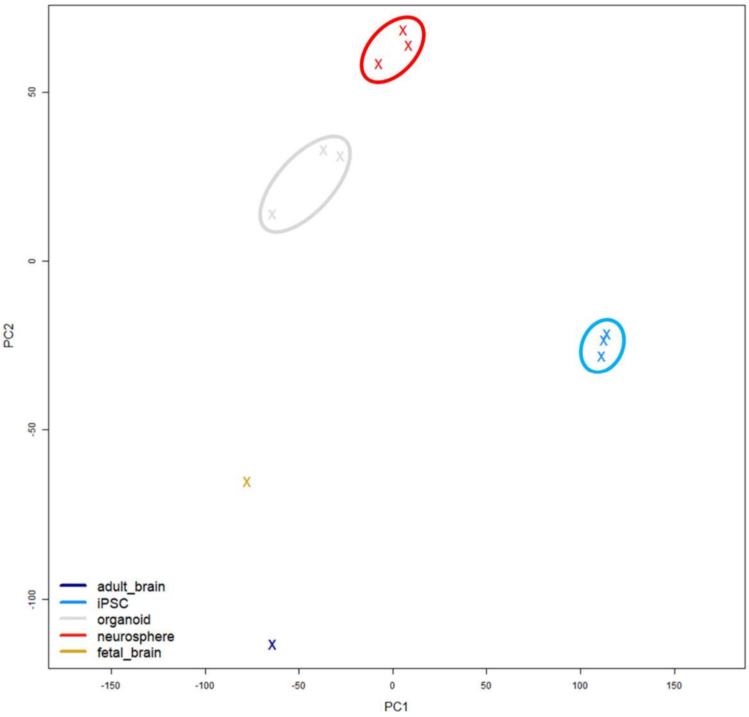


**Supplementary Figure S6: Quality control of next generation sequencing data. (A)** Dendrogram obtained by hierarchical cluster analysis of NGS gene expression data for B4, CS789 and IUFi001 iPSCs, neurospheres and cerebral organoids. (**B**) Principal component analysis of NGS gene expression data for B4, CS789 and IUFi001 iPSCs, neurospheres and cerebral organoids.

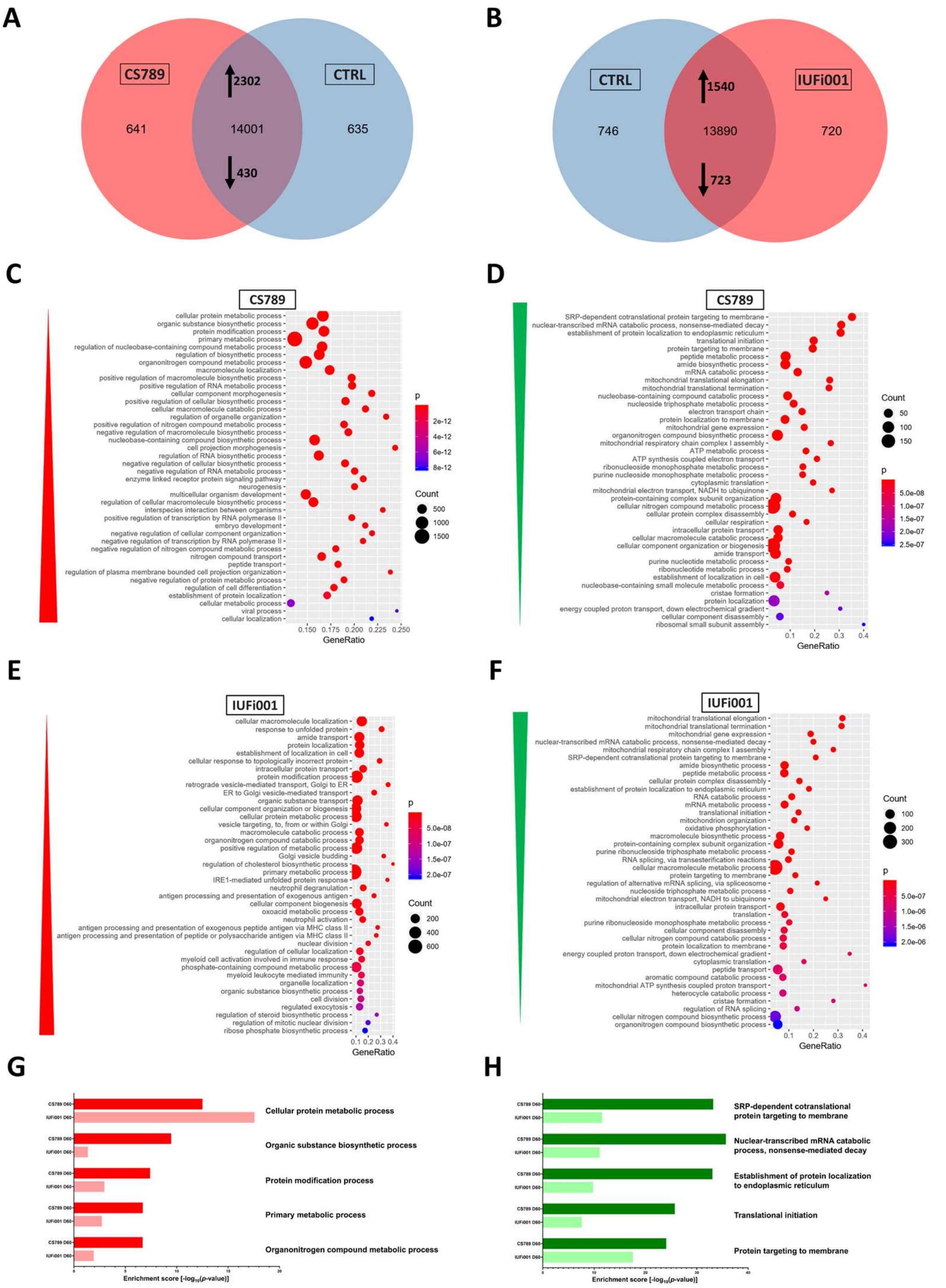


**Supplementary Figure S7: Global transcriptome and associated Gene Ontology analysis of control and CS neurospheres at day 30.** (**A**) Venn diagram showing genes expressed only in CS789 neurospheres (641), in CTRL (B4) neurospheres (635) and common to both (14001) (detection p value < 0.05). (**B**) Venn diagram showing genes expressed only in IUFi001 neurospheres (720), in CTRL (B4) neurospheres (746) and common to both (13890) (detection p value < 0.05). (**C,D**) Dot plots showing the Top 30 differentially regulated Gene Ontologies (C) in the 2302 significantly upregulated DEGs in day 30 CS789 neurospheres in comparison to CTRL (B4) (D) and in the 430 significantly downregulated DEGs in day 30 CS789 neurospheres in comparison to CTRL (B4). (**E,F**) Dot plots showing the Top 30 differentially regulated KEGG pathways (E) in the 1540 significantly upregulated DEGs in day 30 IUFi001 neurospheres in comparison to CTRL (B4) (F) and in the 723 significantly downregulated DEGs in day 30 IUFi001 neurospheres in comparison to CTRL (B4). (**G**) Bar chart of the differentially upregulated Gene Ontologies (Top 5 ranked) common between day 30 CS789 and IUFi001 neurospheres in comparison to CTRL (B4) neurospheres. (**H**) Bar chart of the differentially downregulated Gene Ontologies(Top 5 ranked) common between day 30 CS789 and IUFi001 neurospheres in comparison to CTRL (B4) neurospheres.

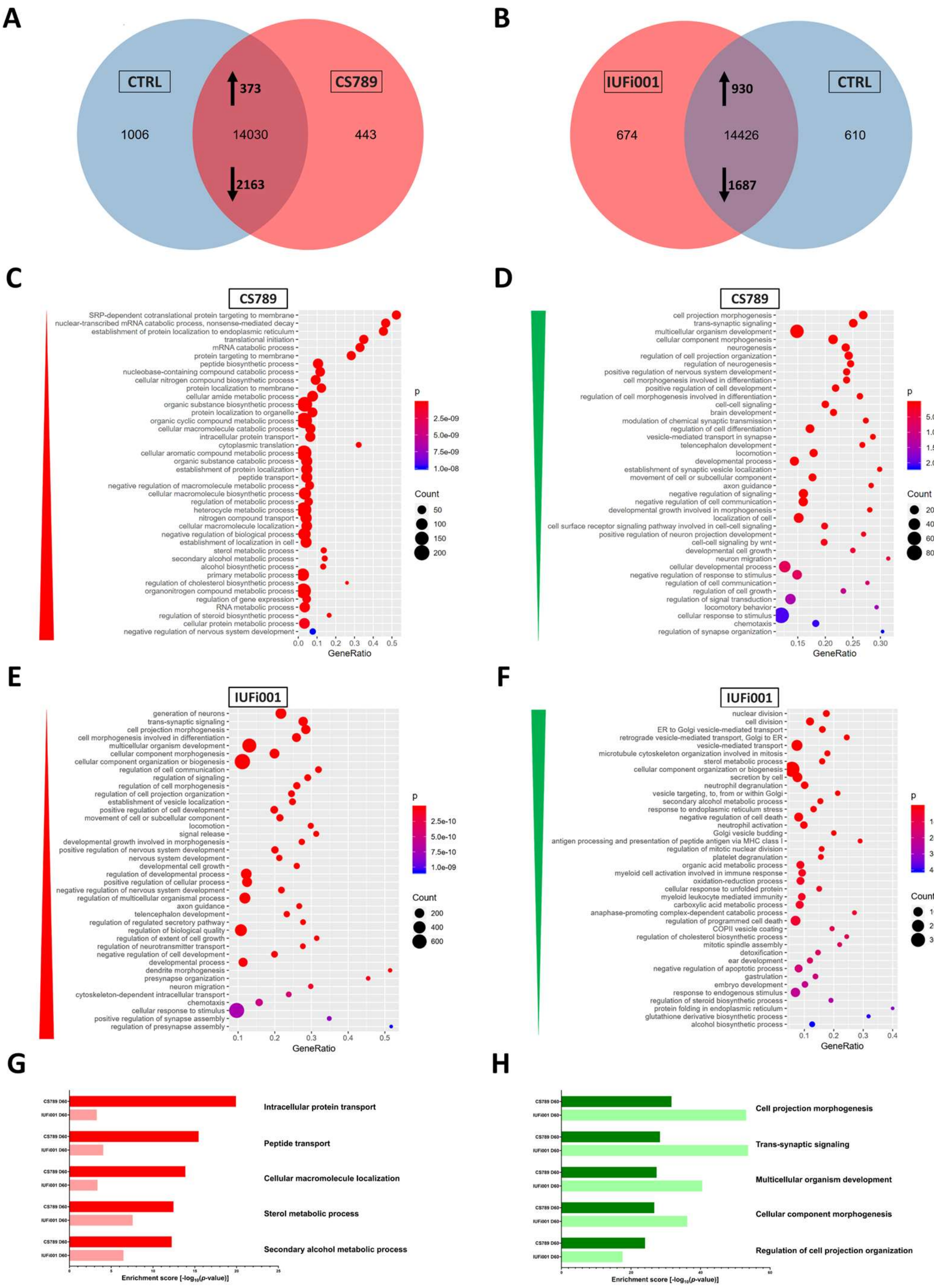


**Supplementary Figure S8: Global transcriptome and associated Gene Ontology analysis of control and CS organoids at day 60.** (**A**) Venn diagram showing genes expressed only in CS789 organoids (443), in CTRL (B4) organoids (1006) and common to both (14030) (detection p-value < 0.05). (**B**) Venn diagram showing genes expressed only in IUFi001 organoids (674), in CTRL (B4) organoids (610) and common to both (14426) (detection p value < 0.05). (**C,D**) Dot plots showing the Top 30 differentially regulated Gene Ontologies (C) in the 373 significantly upregulated DEGs in day 60 CS789 organoids in comparison to CTRL (B4) (D) and in the 2163 significantly downregulated DEGs in day 60 CS789 organoids in comparison to CTRL (B4). (**E,F**) Dot plots showing the Top 30 differentially regulated Gene Ontologies (E) in the 930 significantly upregulated DEGs in day 60 IUFi001 organoids in comparison to CTRL (B4) (F) and in the 1687 significantly downregulated DEGs in day 60 IUFi001 organoids in comparison to CTRL (B4). (**G**) Bar chart of the differentially upregulated Gene Ontologies (Top 5 ranked) common between day 60 CS789 and IUFi001 organoids in comparison to CTRL (B4) organoids. (**H**) Bar chart of the differentially downregulated Gene Ontologies (Top 5 ranked) common between day 60 CS789 and IUFi001 organoids in comparison to CTRL (B4) organoids.

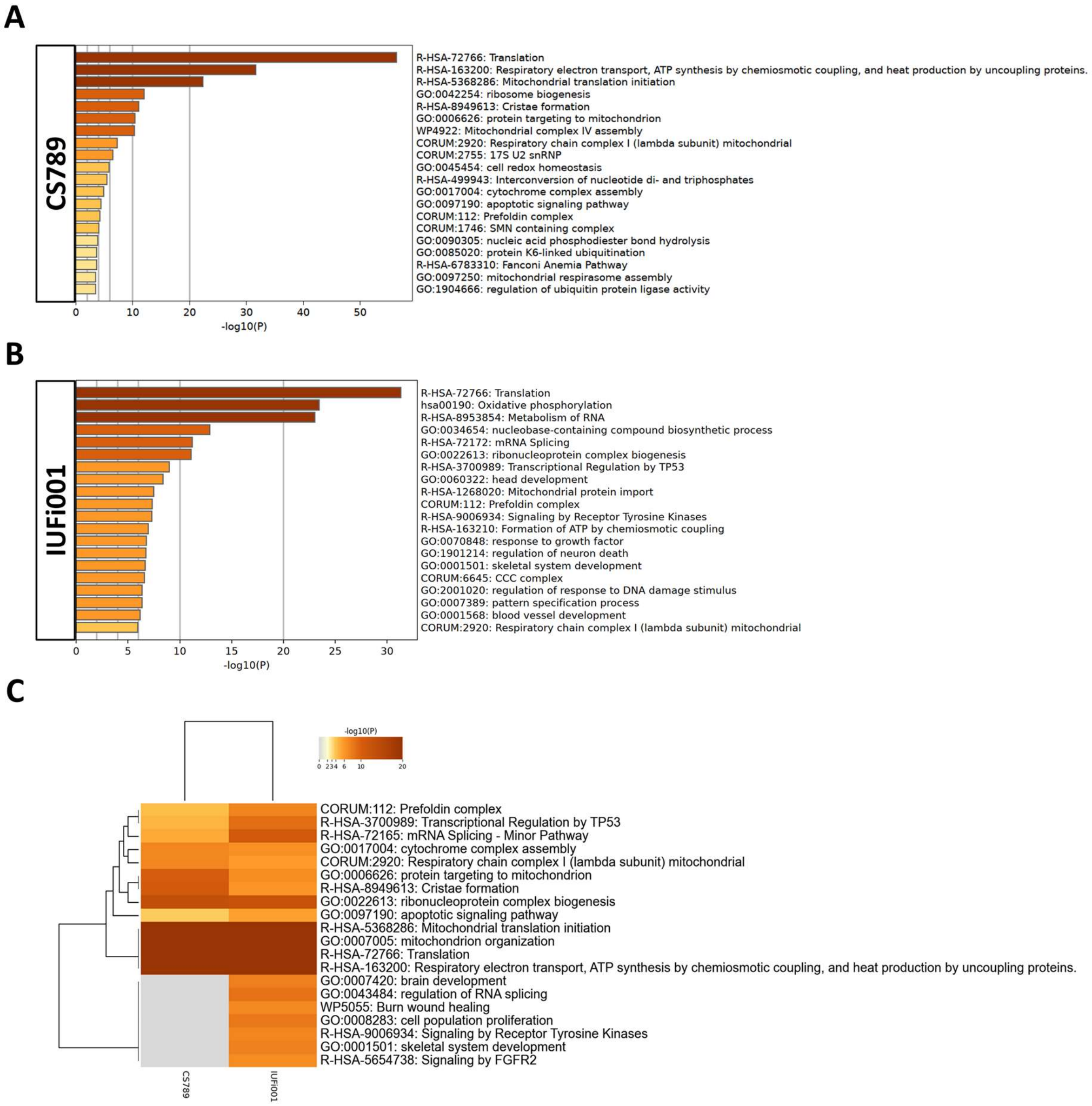


**Supplementary Figure S9: Comparative transcriptome and Gene Ontology analysis of downregulated DEGs in day 30 neurospheres.** (**A**) Bar graph of the Top 20 non-redundant enrichment clusters attributable to the 430 DEGs upregulated in day 30 CS789 neurospheres in comparison to CTRL (B4). (**B**) Bar graph of the Top 20 non-redundant enrichment clusters attributable to the 723 DEGs downregulated in day 30 IUFi001 neurospheres. (**C**) Metascape-generated heatmap comparing downregulated gene-sets employed in A and B revealed i.a. GOs involved in mitochondrial translation initiation, mitochondrion organization and translation.

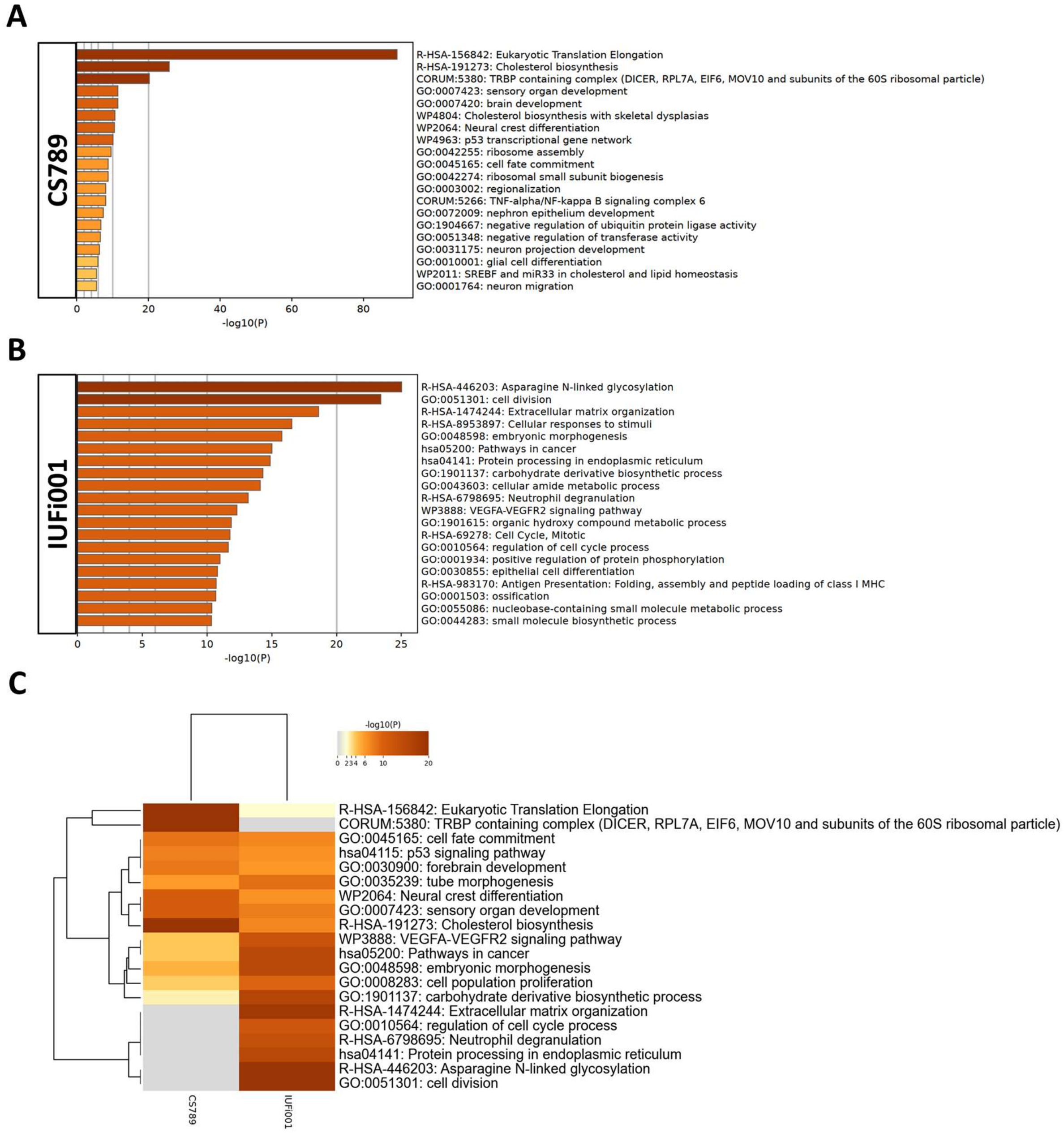


**Supplementary Figure S10: Comparative transcriptome and Gene Ontology analysis of upregulated DEGs in day 60 organoids.** (**A**) Bar graph of the Top 20 non-redundant enrichment clusters attributable to the 373 DEGs upregulated in day 60 CS789 (D1) organoids in comparison to CTRL (B4). (**B**) Bar graph of the Top 20 non-redundant enrichment clusters attributable to the 930 DEGs upregulated in day 60 IUFi001 organoids. (**C**) Metascape-generated heatmap comparing upregulated gene-sets employed in A and B revealed i.a. GOs involved in forebrain development, p53-signalling pathway, cholesterol biosynthesis and VEGFA-VEGFR2 signalling pathway.

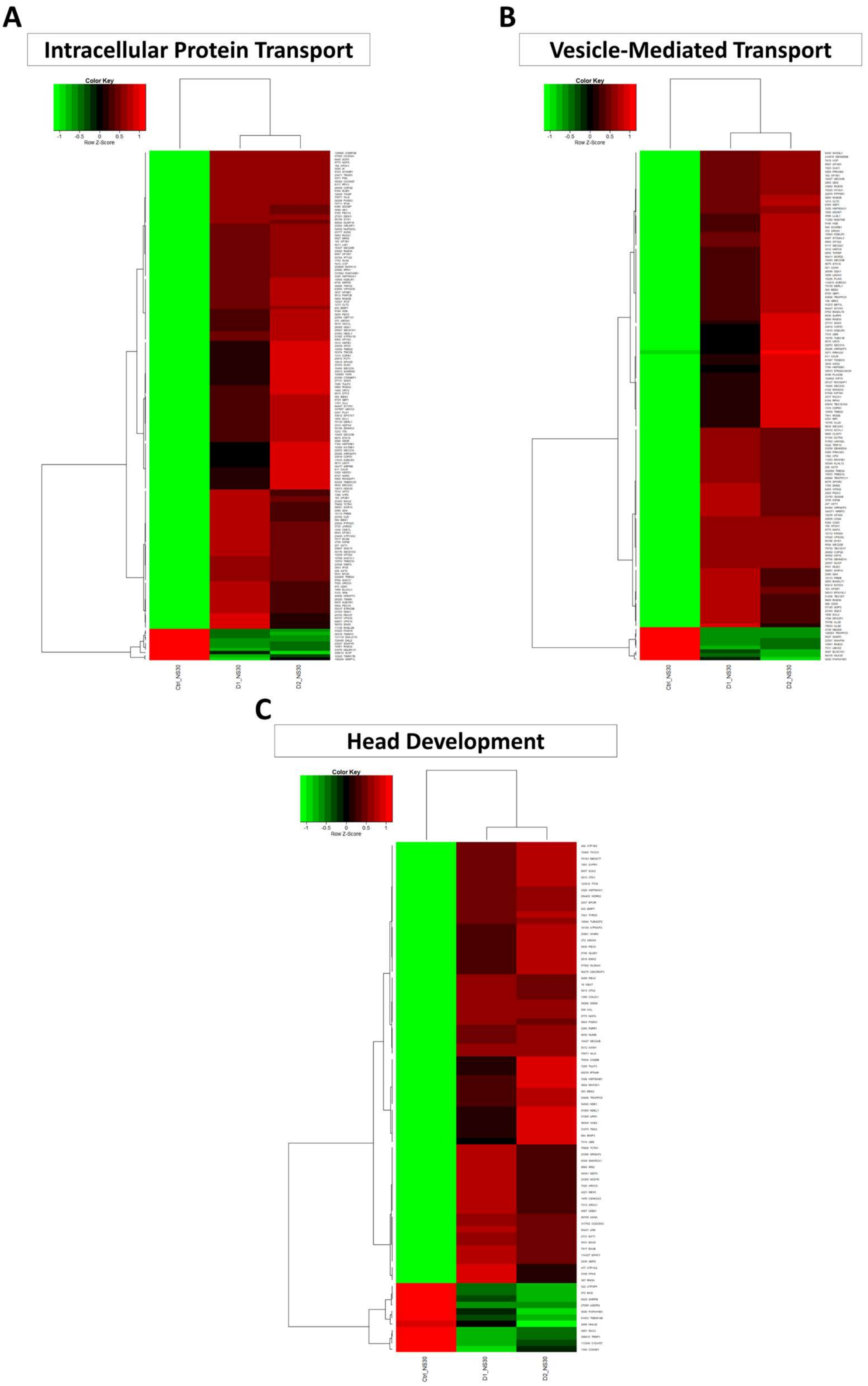


**Supplementary Figure S11: Analysis of select Gene Ontologies differentially regulated in day 30 neurospheres.** (**A**) Pearson´s heatmap depicting all identified differentially regulated genes involved in intracellular protein transport common between day 30 CS789 (D1) and IUFi001 (D2) neurospheres in comparison to B4 (CTRL). (**B**) Pearson´s heatmap depicting all identified differentially regulated genes involved in vesicle-mediated transport common between day 30 CS789 (D1) and IUFi001 (D2) neurospheres in comparison to B4 (CTRL). (**C**) Pearson´s heatmap depicting all identified differentially regulated genes involved in head development common between day 30 CS789 (D1) and IUFi001 (D2) neurospheres in comparison to B4 (CTRL).

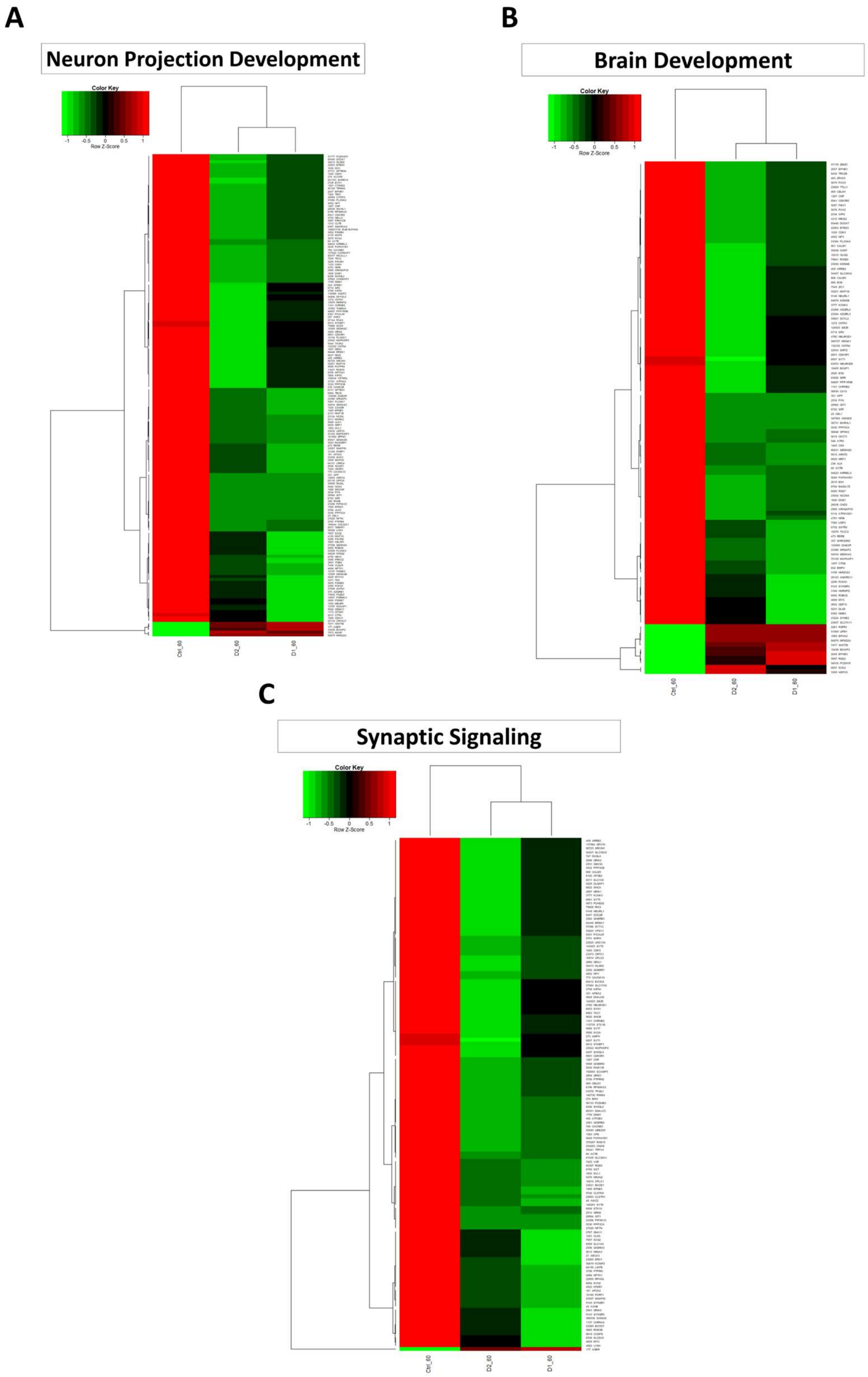


**Supplementary Figure S12: Analysis of select Gene Ontologies differentially regulated in day 60 organoids.** (**A**) Pearson´s heatmap depicting all identified differentially regulated genes involved in neuron projection development common between day 60 CS789 (D1) and IUFi001 (D2) organoids in comparison to B4 (CTRL). (**B**) Pearson´s heatmap depicting all identified differentially regulated genes involved in brain development common between day 60 CS789 (D1) and IUFi001 (D2) organoids in comparison to B4 (CTRL). (**C**) Pearson´s heatmap depicting all identified differentially regulated genes involved in synaptic signalling common between day 60 CS789 (D1) and IUFi001 (D2) organoids in comparison to B4 (CTRL).

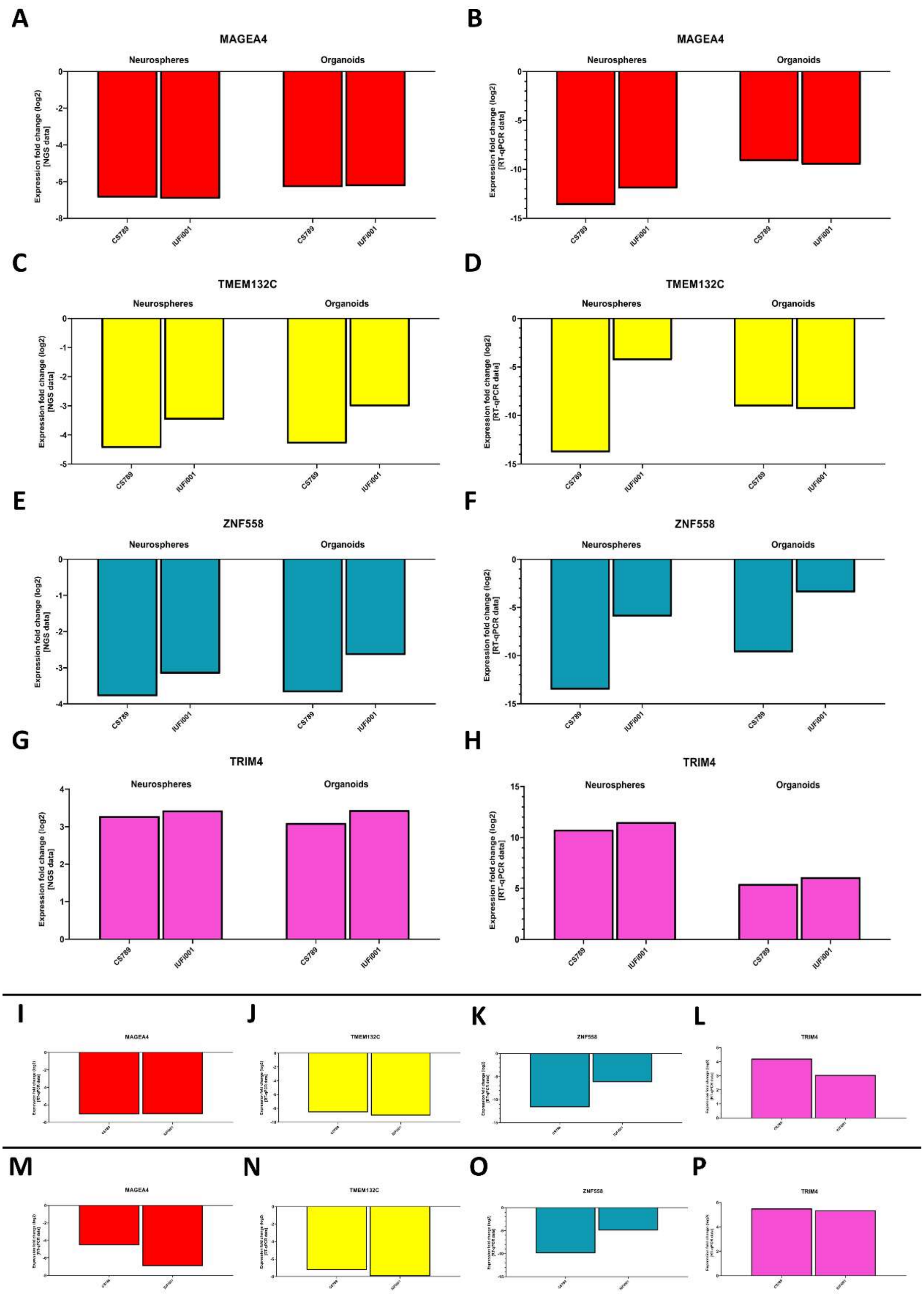


**Supplementary Figure S13: RT-qPCR for MAGEA4, TMEM132C, ZNF558 and TRIM4 in Day 30 Neurospheres, two sets of Day 60 Organoids and Day 120 Organoids.** (**A, C, E, G**) Relative mRNA expression analysis of *MAGEA4* (A), *TMEM132C* (C), *ZNF558* (E), and *TRIM4* (G) in CS789 and IUFi001 day 30 neurospheres and day 60 organoids compared to CTRL (B4). (**B, D, F, H**) qRT-PCR analysis of *MAGEA4* (B), *TMEM132C* (D), *ZNF558* (F), and *TRIM4* (H) mRNA expression in CS789 and IUFi001 day 30 neurospheres and day 60 organoids relative to CTRL (B4). (**I-L**) RT-qPCR analysis of *MAGEA4* (I), *TMEM132C* (J), *ZNF558* (K) and *TRIM4* (L) mRNA expression in a second set of day 60 CS789 and IUFi001 organoids relative to control organoids (B4). **(M-P)** RT-qPCR analysis of *MAGEA4* (M), *TMEM132C* (N), *ZNF558* (O) and *TRIM4* (P) mRNA expression of day 120 CS789 and IUFi001 organoids relative to control organoids (B4).

**A**

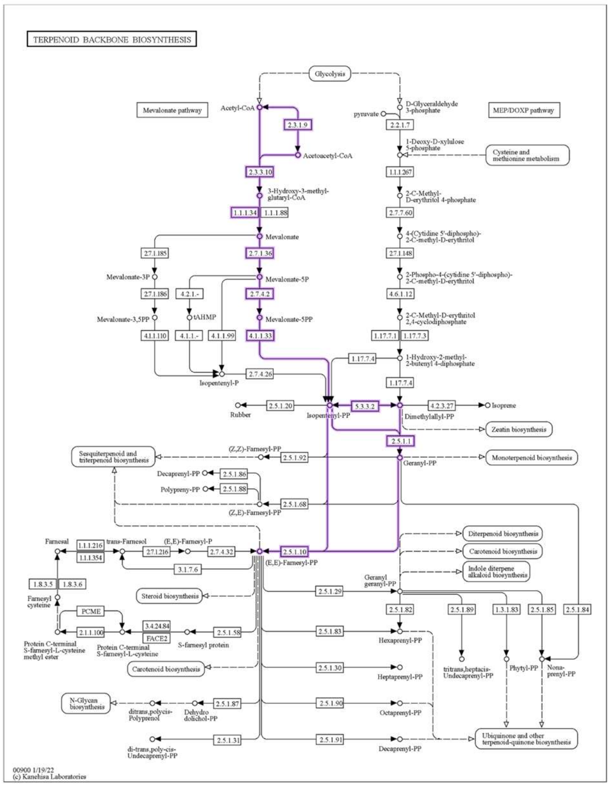


**B**

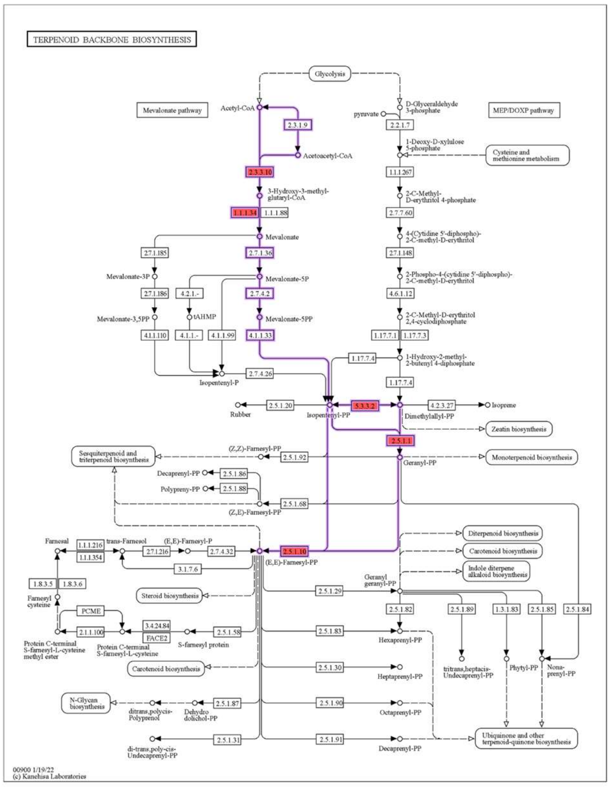


**Supplementary Figure S14: Analysis of the Mevalonate pathway of the Terpenoid Backbone Biosynthesis pathway in day 30 NS and day 60 COs.** (**A, B**) Schematic of the pathway Terpenoid Back Biosynthesis with the Mevalonate pathway indicated in purple. The genes upregulated in CSB-deficient day 30 NS (A) and day 60 COs (B) are indicated in red.

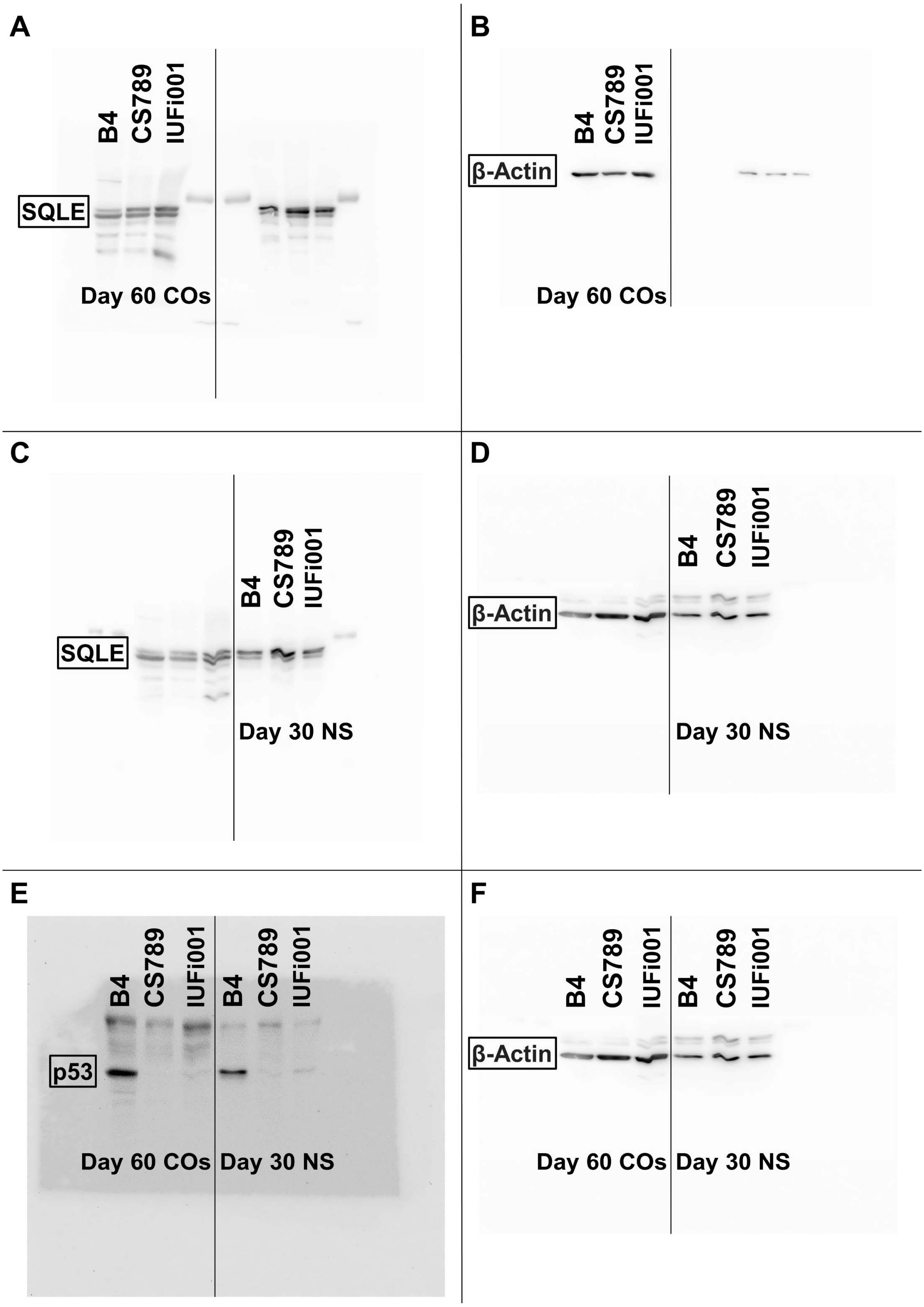


**Supplementary Figure S 15: Full Western Blot for CSB, p53 and β-Actin protein in Day 30 NS and Day 60 COs.** (**A**) Western Blot analysis for SQLE protein in B4, CS789 and IUFi001 COs. (**B**) Western Blot for beta-Actin protein in B4, CS789 and IUFi001 COs. (**C**) Western Blot analysis for SQLE protein in B4, CS789 and IUFi001 NS. (**D**) Western Blot for beta-Actin protein in B4, CS789 and IUFi001 NS. (**E**) Western Blot analysis for p53 protein in B4, CS789 and IUFi001 NS and COs. (**F**) Western Blot for beta-Actin protein in B4, CS789 and IUFi001 NS and COs. (**B, D, F**) Blot of the housekeeping gene used to quantify the respective blot in A, C and E on the left side of the figure.
